# Systematic Functional Interrogation of Neurodevelopmental Disorder Risk Variants

**DOI:** 10.64898/2026.10.21.760732

**Authors:** Alejandro Gomez, Jessica C. McAfee, Hyunggyu Min, Tatiana Ulloa Avila, Sean Jones, Jessica L. Bell, Yangzhenyu Gao, Ellen Hu, Haoyue Li, Sool Lee, Ratchell Sadovnik, Zheng Fan, Nan Yang, Adriana S. Beltran, Hyejung Won

## Abstract

Neurodevelopmental disorders (NDDs) are highly polygenic, yet the functional consequences of most associated non-coding variants remain unclear. We used a massively parallel reporter assay (MPRA) to test 727 NDD-associated non-coding variants in human neural progenitors, identifying 38 expression-modulating variants (emVars) with regulatory activity. These emVars were predicted to alter transcription factor binding motifs linked to neurodevelopmental signaling and cellular metabolism. We mapped emVars to candidate target genes using expression quantitative trait loci and activity-by-contact (ABC) models. We then used CRISPR droplet sequencing (CROP-seq) in human pluripotent stem cell-derived neurons to identify endogenous target genes and characterize their downstream transcriptional effects. CROP-seq preferentially supported ABC predictions, identified additional regulatory relationships, and revealed convergence on developmentally regulated programs involving neuronal connectivity, synaptic function, and mitochondrial biology. Together, these results connect NDD risk variants to endogenous target genes and shared neurodevelopmental transcriptional programs.

## INTRODUCTION

Neurodevelopmental disorders (NDDs), including autism spectrum disorder (ASD) and attention deficit hyperactivity disorder (ADHD), comprise a heterogeneous group of conditions characterized by early-onset impairments in cognition, behavior, and social functioning. Collectively, NDDs affect millions of individuals worldwide and impose substantial societal and economic burdens. Genetic studies of NDDs have identified rare, highly penetrant mutations with large effect sizes, including *de novo* protein-truncating variants^1–4^ and copy number variants^5–7^, which have provided important insights into biological pathways underlying NDD risk. However, accumulating evidence suggests that common variants of individually modest effect collectively account for a substantial proportion of the heritable liability underlying NDDs.

Genome-wide association studies (GWAS) of ASD^8,9^, ADHD^10,11^, and a cross-disorder meta-analysis of multiple NDDs^12^ (hereafter referred to as pan-NDD) have begun to identify common genetic variants associated with NDD susceptibility. However, the vast majority of GWAS variants reside in non-coding regions of the genome and are inherited in linkage disequilibrium (LD) with dozens of neighboring variants, complicating efforts to identify causal variants and define their downstream molecular consequences. Systematic functional characterization of non-coding risk variants is therefore essential for elucidating mechanisms through which common genetic variants contribute to NDD pathogenesis.

Given that clinical manifestations of NDDs typically emerge during early childhood, understanding how genetic risk factors perturb transcriptional programs during neurodevelopment is likely critical for elucidating NDD etiology. Early developmental processes, including neural progenitor proliferation, neuronal differentiation, and neural circuit assembly, are orchestrated by tightly regulated transcriptional programs that may be particularly vulnerable to disruption by regulatory variation. Defining how non-coding risk variants influence these developmental transcriptional networks may therefore advance our understanding of NDD etiology.

Here, we combined a massively parallel reporter assay (MPRA), integrative variant-gene mapping strategies, and CRISPR-based perturbation to systematically characterize the molecular consequences of NDD-associated variants. First, we employed MPRA to directly quantify the regulatory activity of hundreds of NDD-associated variants in human neural progenitor cells (NPCs), a cellular model of early cortical development^13,14^. We then integrated expression quantitative trait loci (eQTL) as well as activity-by-contact (ABC) models to nominate putative target genes for regulatory variants identified by MPRA. To experimentally evaluate the predicted variant-gene relationships, we performed CRISPR droplet sequencing (CROP-seq). Finally, we delineated transcriptional networks perturbed by NDD-associated variants and demonstrated that common variants, despite their individually modest effects, can collectively elicit coordinated transcriptional changes that converge on key neurodevelopmental processes.

Together, our integrative strategy closes the loop from genetic association to regulatory function, target gene identification, and downstream transcriptional consequences. These findings advance our understanding of the regulatory landscape underlying NDD risk and provide a roadmap for future efforts to connect human genetic variation with disease-associated molecular pathways.

## RESULTS

### Identification of NDD-associated functional regulatory variants

We compiled 18 genome-wide significant (GWS) loci from three NDD GWAS datasets: ASD^8,9^, ADHD^10^, and pan-NDD^12^ (**Figure 1A**). To construct an MPRA library, we selected 727 variants that (1) were located within GWS loci, (2) had a GWAS association of P < 1×10^−5^, and (3) did not contain restriction enzyme recognition sites used for downstream molecular cloning (**Figure 1A-B**, **Methods**). Of the selected variants, 133 were from the ASD GWAS, 479 from the ADHD GWAS, and 184 from the pan-NDD GWAS, with 69 variants shared between ADHD and pan-NDD (**Figure 1A**). We also included 6 positive control sequences consisting of 3 tiled fragments from the cytomegalovirus (CMV) and the eukaryotic translation elongation factor 1 alpha (EF1α) promoters, as well as 200 scrambled negative control sequences. The resulting NDD-MPRA library comprised a 150-base-pair (bp) genomic sequence centered on the variant, followed by a minimal promoter, a luciferase gene, and a 20-bp barcode uniquely mapped to each variant (**Figure 1B**).

**Figure 1.**
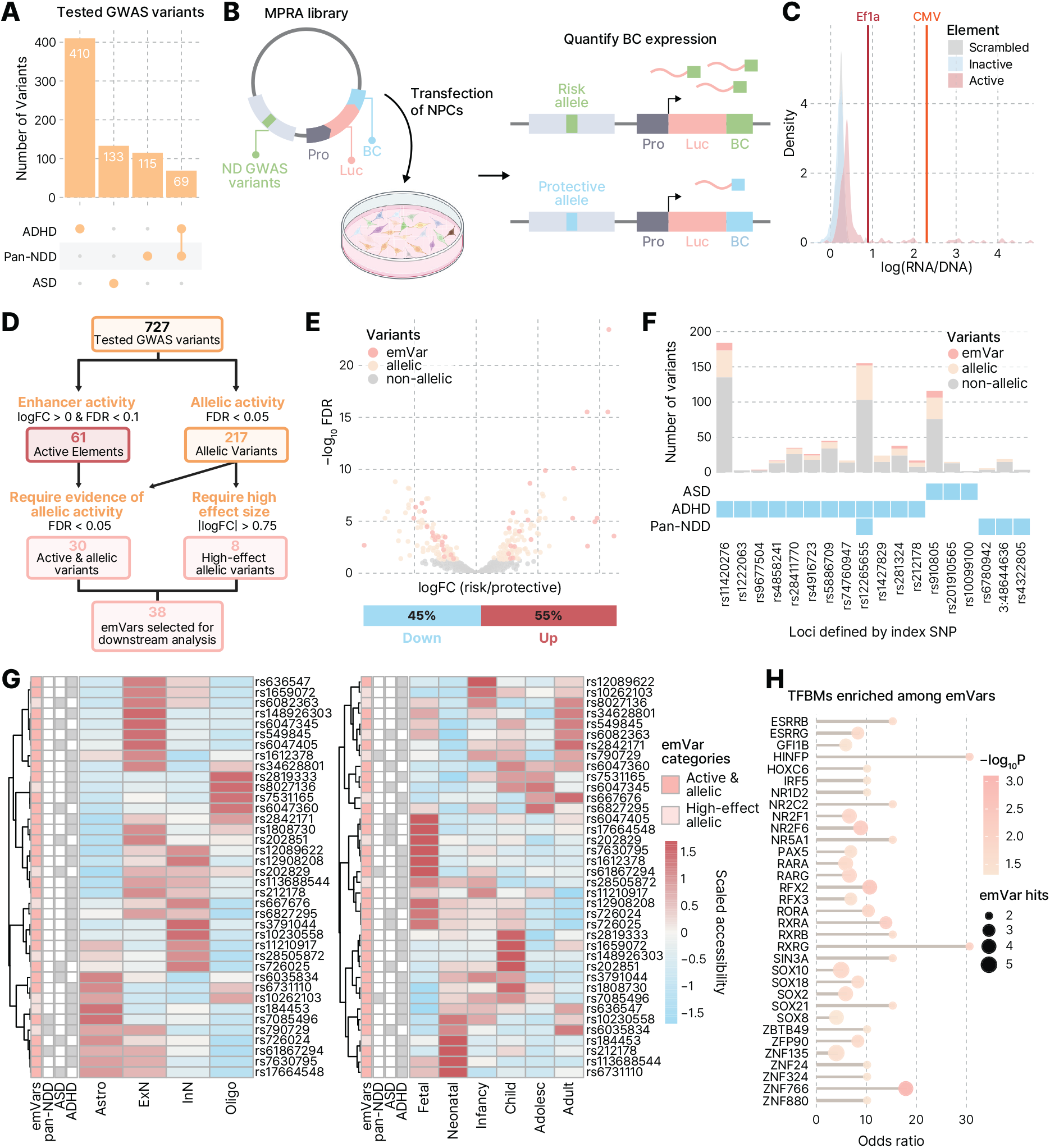
Characterization of NDD-associated MPRA variants. **A**. Number of GWAS variants selected for MPRA testing from ASD, ADHD, and pan-NDD loci. **B**. Schematic of the NDD-MPRA library. Each tested element contained a 150-bp genomic sequence centered on an NDD risk variant, followed by a minimal promoter (Pro), luciferase reporter, and uniquely assigned barcode (BC). The resulting NDD-MPRA library was introduced to NPCs, where regulatory differences between risk and protective alleles were quantified by barcode expression. **C**. Density plot of mean log(RNA/DNA) ratios for tested elements and scrambled negative controls. Positive controls, including EF1a and CMV promoter sequences, are indicated by vertical lines. **D**. Schematic of the variant prioritization strategy used to define emVars. **E.** Volcano plot showing allelic regulatory activity across MPRA-tested variants, with dots colored by MPRA regulatory class. The x-axis represents MPRA logFC of the risk allele relative to the protective allele. The bar plot below summarizes the direction of risk-allele effects among emVars. **F**. Number of variants tested per locus, defined by the index SNP and colored by MPRA regulatory class. Blue boxes indicate the disorder association of each locus. **G**. Scaled chromatin accessibility of emVars across major brain cell types (left) and across developmental stages of excitatory neurons (right). Rows represent individual emVars, with annotations indicating emVar class and disorder association. **H**. Nominal enrichment of TFBMs among emVars compared with matched background SNPs, measured by one-sided Fisher’s exact test. Dot size indicates the number of emVars associated with each TF motif, odds ratio indicates the magnitude of enrichment relative to background SNPs, and color indicates –log10P.

Because genetic risk factors for NDDs are enriched in regulatory elements of the developing brain^8,12,15,16^, we conducted MPRA in NPCs, as previously described^13,14^. Following transfection of the NDD-MPRA library into NPCs, barcode DNA and RNA were extracted and processed for high-throughput sequencing (**Figure 1B**, **Methods**). The assay showed high reproducibility across 22 biological replicates, with a median pairwise Pearson correlation of r = 0.897 across 231 pairwise comparisons (**Figure S1A**).

For each variant, we first assessed element-level MPRA activity by comparing the RNA/DNA ratio of each variant-containing element with scrambled negative controls and classified NDD variants as inactive and active elements [active false discovery rate (FDR) < 0.1, **Figure S1B**]. Consistent with this classification, scrambled controls and inactive NDD elements showed similar log(RNA/DNA) distributions, whereas active NDD elements exhibited higher reporter activity. CMV and EF1α positive control sequences showed the highest reporter activity, further supporting the distinction of active and inactive elements (**Figure 1C**).

We then assessed allelic regulatory activity by comparing RNA/DNA ratios between the risk and protective alleles of each variant (**Figure S1C**). Among the 727 variants that passed quality control (QC), 479 showed neither significant allelic effects nor significant element-level activity, 179 variants showed significant allelic effects only (allelic FDR < 0.05), and 31 showed significant element-level activity only (active FDR < 0.1). We identified 30 variants that exhibited both significant allelic effects and element-level activity. We further included 8 additional variants that lacked significant element-level activity but showed strong allelic effects (|allelic logFC| > 0.75; **Figure S1B-C**). Together, 38 variants (30 with both allelic and element-level activity and 8 with strong allelic effects) were defined as expression-modulating variants (emVars) and carried forward for downstream analyses (**Figure S1D**, **Figure 1D-E**, **Table S1**). For comparison, we defined background single-nucleotide polymorphisms (SNPs) by matching minor allele frequency and LD, as previously described^13^. The directionality of risk-allele effects among emVars was balanced, with 17/38 (44.7%) showing lower and 21/38 (55.3%) showing higher reporter activity relative to the protective allele, suggesting no strong systematic directional bias in the effects of risk alleles on regulatory activity (**Figure 1E**, **bottom**).

Of the 18 tested loci, 10 contained at least one emVar. Five loci contained allelic variants but no emVars, whereas three loci lacked both MPRA-allelic variants and emVars, although these loci contained only a small number of tested variants (2-4 variants per locus). In total, the 38 emVars comprised 10 ASD-associated variants, 25 ADHD-associated variants, and 3 variants shared between ADHD and pan-NDD (**Figure 1F**).

### Characteristics of NDD emVars

We next compared emVars with endogenous chromatin states annotated by chromHMM^17^ (**Figure S2A**). Compared with background SNPs, emVars showed greater representation in both active and repressive chromatin states in human pluripotent stem cell (hPSC)-derived neurons (H9 neurons), neurospheres, and the fetal brain (**Figure S2B**). The overall fraction of emVars overlapping any annotated chromatin state was higher in the fetal brain compared to the adult brain (**Figure S2B**), consistent with stronger engagement of NDD emVars within developmentally regulated chromatin contexts.

We previously showed that MPRA-validated enhancer variants associated with psychiatric disorders and human cortical structure are enriched for Alu elements^14,18^, which can regulate gene expression and have been implicated in human brain evolution^19^. We therefore examined whether NDD emVars showed a similar relationship with repetitive elements. Compared with matched background SNPs, emVars showed substantially greater overlap with Alu elements (47% versus 14%, FDR = 4.8×10^−6^; **Figure S2C**). In contrast, no significant enrichment was observed for other classes of repetitive elements such as SVA, L1, LTR, and hAT-Charlie elements (**Figure S2C**). Together with previous findings, these results support a potential contribution of repetitive elements to the regulatory activity of neurodevelopmental and psychiatric disorder-associated variants.

To further characterize the cellular and developmental contexts of NDD emVars, we examined chromatin accessibility across major brain cell types and developmental stages of excitatory neurons using a human brain single-cell atlas^20^. Accessibility profiles varied across emVars, although 24 of 38 (63.2%) showed their highest chromatin accessibility in excitatory or inhibitory neurons among the cell types examined (**Figure 1G**). This is consistent with our previous observations that emVars in NPCs can retain regulatory activity in postmitotic neuronal contexts^13,14^. Across developmental stages of excitatory neurons, 27 of 38 emVars (71.1%) showed their highest accessibility by childhood, supporting a developmental bias in the chromatin contexts harboring NDD emVars.

We next sought to identify transcriptional regulators through which emVars may influence gene expression. Using MotifbreakR^21,22^, we identified transcription factor binding motifs (TFBMs) predicted to be created or disrupted by risk alleles (**Figure 1H**, **Figure S2D**, **Table S2**). We identified 33 nominally enriched TFBMs with predicted differential binding between risk and protective alleles compared with matched background SNPs (one-sided Fisher’s exact test, P < 0.05; **Methods**). Several implicated transcription factors (TFs) showed distinct expression patterns across neural differentiation. Progenitor- and lineage-associated factors, including SOX2, SOX8, SOX10, and SOX21, showed relatively higher expression in NPCs, whereas nuclear receptor-related factors, including RORA, RARA, ESRRB, and ESRRG, showed relatively higher expression in neurons (**Figure S2E**). These TFs encompass regulators of neural progenitor maintenance and neuronal differentiation^23–27^, as well as transcriptional programs involved in mitochondrial and metabolic function^28^. Notably, disruption of several of these transcriptional regulators has also been linked to NDDs. Pathogenic *SOX2* variants are associated with developmental disability^29,30^, whereas *NR2F1* haploinsufficiency causes Bosch-Boonstra-Schaaf Optic Atrophy Syndrome^31^. Pathogenic *RORA* variants have been similarly implicated in syndromic NDDs characterized by intellectual disability, autism, and cerebellar phenotypes^32,33^. Together, these findings suggest that NDD emVars may alter regulatory sites used by TF networks that coordinate neural differentiation and energy regulation.

### Integrative variant-to-gene mapping prioritizes genes implicated in neurodevelopment

Given that emVars are expected to influence disease risk through gene regulation, we next used two complementary approaches, eQTL and ABC models, to nominate putative target genes for the 38 emVars (**Figure 2A**, **Methods**). For eQTL mapping, we used resources from two developmental epochs—fetal^34^ and adult^35^ brain—to assess whether the putative target genes of NDD emVars vary across brain development. Because both adult and fetal brain eQTL datasets were generated from bulk brain tissue and therefore lack cell type specificity, we additionally incorporated an adult postmortem brain single-nucleus eQTL (sn-eQTL) resource^36^ that provides cell type-resolved regulatory relationships (**Figure 2A**). We identified putative target genes for emVars across neuronal development by generating high-resolution chromatin contact maps from human induced pluripotent stem cell (hiPSC)-derived NPCs and neurons using Micro-C (**Methods**) and integrating these maps with corresponding chromatin accessibility data^37,38^ (**Figure 2A**).

**Figure 2.**
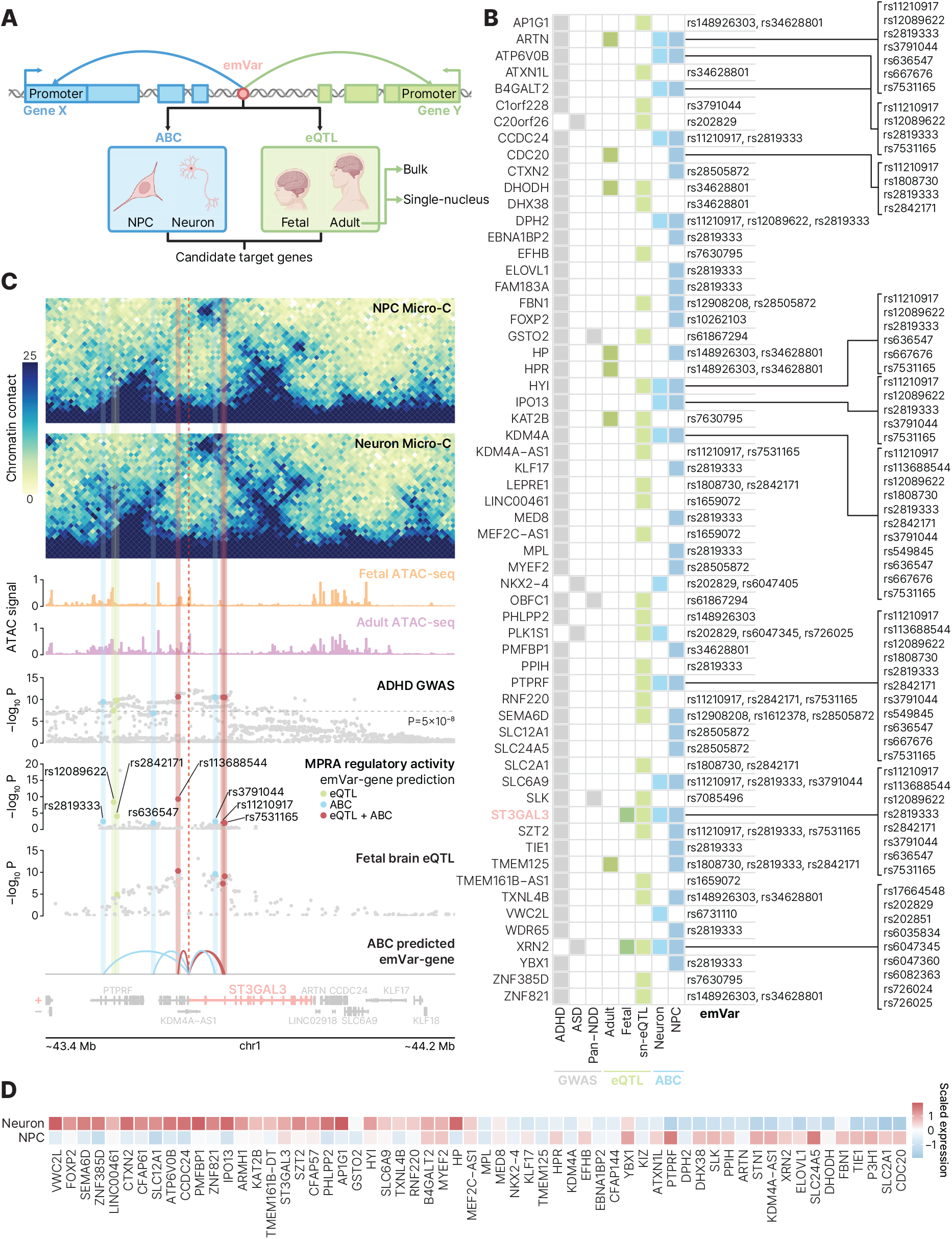
Mapping emVars to target genes. **A.** Schematic of emVar-gene mapping methods. The pink circle represents an emVar. emVars were mapped to putative target genes by ABC (blue), which leverages cell type-specific chromatin accessibility and Micro-C data from NPCs and neurons, and eQTL (green), which leverage fetal brain (Fetal) and adult brain (Adult) bulk RNA-seq or snRNA-seq (sn-eQTL) datasets. **B.** Summary of genes mapped to emVars by different gene mapping methods. Rows represent mapped genes, and columns indicate the associated disorder, emVar-gene mapping method, and linked emVar(s). eQTL-based gene mappings are denoted in green, whereas ABC-based predictions are denoted in blue. *ST3GAL3*, highlighted in red, is further characterized in (C). **C**. *ST3GAL3* locus plot with normalized Micro-C contact profiles for NPCs and neurons, fetal and adult neuronal ATAC-seq signals, ADHD GWAS association statistics, MPRA regulatory activity, fetal brain eQTL association statistics, and gene annotations. emVars mapped within this locus are colored by their gene mapping methods: eQTL only (green), ABC only (blue) and both eQTL and ABC (red). Semi-transparent vertical lines indicate emVar positions, and the dashed red line denotes the *ST3GAL3* promoter. **D**. Expression of emVar-linked genes across neuronal differentiation stages from the LIBD Stem Cell Browser. Gene-wise z scores were calculated across five differentiation conditions and subsequently restricted to NPCs and neurons for comparison. The complete heatmap across all differentiation states is shown in Figure S3D.

Among the 38 emVars, these approaches nominated 144 unique variant-gene pairs involving 32 emVars and 60 genes (**Figure 2B**, **Table S3**). Collectively, the eQTL datasets contributed 66 variant-gene pairs across 37 genes, whereas ABC models contributed 87 pairs across 37 genes, with 9 pairs being supported by both approaches. Among individual resources, NPC-ABC produced the largest number of predictions, identifying 59 pairs involving 34 genes, followed by sn-eQTL (54 pairs involving 32 genes), neuron-ABC (39 pairs involving 15 genes), adult brain eQTL (12 pairs involving 7 genes), and fetal brain eQTL (6 pairs across 2 genes) (**Figure S3A**).

NPC- and neuron-ABC showed relatively greater concordance than the eQTL resources, although NPC-ABC nominated more target genes than neuron-ABC (**Figure S3B**). This concordance suggests that a subset of predicted variant-gene relationships is maintained across neuronal differentiation, while the larger number of NPC-ABC predictions may reflect the regulatory context in which the emVars were originally identified. In contrast, fetal and adult brain eQTL analyses identified largely distinct sets of target genes, consistent with developmental changes in cellular composition and gene regulatory programs. sn-eQTL identified more target genes than either bulk fetal or adult brain eQTL, consistent with the strong cell type-specificity of genotype-expression relationships. Across all mapping approaches, most nominated genes were linked to 1-2 emVars, while a subset of emVars was linked to multiple candidate genes, indicating that emVar-gene relationships are not uniformly one-to-one (**Figure S3C**).

Despite limited concordance between eQTL and ABC predictions^13,14^, several genes showed convergent support across orthogonal datasets, including 6 genes (*HYI*, *ARTN*, *PTPRF*, *KDM4A*, *XRN2*, and *ST3GAL3*) that were nominated by at least 3 independent resources. Several have established roles in neurodevelopmental processes. *ARTN* encodes a neurotrophic factor that regulates neuronal migration, neurite outgrowth, and neuronal survival^39,40^. *KDM4A* encodes a histone demethylase involved in neural lineage specification and differentiation^41,42^. *PTPRF* encodes a member of the LAR receptor protein tyrosine phosphatase family, which regulates axon development and synaptic organization^43,44^. *XRN2* encodes a nuclear 5′–3′ exoribonuclease involved in RNA turnover and transcription termination and has been implicated in clearance of developmental transcripts during early stem cell differentiation^45^ and resolution of transcription-associated R-loops^46^. Importantly, several of these repeatedly nominated genes have also been independently implicated in ASD and ADHD genetics. *HYI, PTPRF, and KDM4A* have been identified through gene-based association analyses of ADHD^10^, with *PTPRF* additionally nominated as a putatively causal ADHD gene through Mendelian randomization integrating ADHD genetics with cortical molecular data^47^. ADHD risk alleles have also been associated with increased brain expression of *ARTN*, providing functional evidence linking ADHD-associated variation to regulation of this gene^48^. In ASD, *XRN2* has been implicated by transcriptome-wide association analysis, which found that higher *XRN2* expression was associated with increased ASD risk^49^. Together, these findings indicate that convergence across independent emVar-gene mapping approaches prioritizes genes supported by both neurodevelopmental biology and human NDD genetics.

Among all predicted target genes, *ST3GAL3* emerged as the most strongly supported candidate gene (**Figure 2C**). *ST3GAL3* encodes a sialyltransferase that catalyzes the addition of sialic acid to glycoproteins and glycolipids, a modification with important roles in neural development, synaptic function, and cell-cell interactions^50,51^. Consistent with these functions, pathogenic loss-of-function variants in *ST3GAL3* impair cognitive development and cause *ST3GAL3*-related developmental and epileptic encephalopathy^52,53^. *ST3GAL3* was linked to eight emVars through eQTL, ABC, or both (**Figure 2C**). Notably, the convergence on *ST3GAL3* was not driven by a single variant-gene relationship. Instead, distinct emVars were implicated by different mapping approaches, with 2 emVars linked to *ST3GAL3* exclusively by eQTL, 3 exclusively by ABC, and 3 supported by both approaches. These findings suggest that regulatory variation within the *ST3GAL3* locus may be distributed across multiple functional elements that converge on a common effector gene across neurodevelopmental stages.

Since different variant-gene mapping strategies nominated distinct sets of target genes, we sought to experimentally test emVar-gene relationships using CROP-seq, which combines CRISPR-mediated perturbation with droplet-based single-cell RNA-sequencing (scRNA-seq). To determine whether predicted emVar targets were expressed in cellular contexts relevant to CROP-seq validation, we examined their expression across stem cell-derived neural differentiation states^54^. Among the 60 mapped genes in the expression matrix, 31 showed higher mean expression in differentiated neurons than in NPCs (**Figure 2D, Figure S3D**). When stratified by mapping evidence, genes nominated exclusively by ABC, exclusively by eQTL, or by both approaches spanned a broad range of expression in both NPCs and neurons (**Figure S3E**). Genes supported by both eQTL and ABC showed a higher median expression, although all three groups showed substantial overlap in expression ranges, and a modest shift toward higher expression in neurons (**Figure S3E**). This neuronal shift remained evident when additional differentiation states were considered (**Figure S3D**). Together, these results suggest that emVar-linked genes are expressed across neuronal differentiation, with a modest bias toward higher expression in differentiated neurons, motivating our use of differentiated neurons for CROP-seq.

### CROP-seq identifies endogenous gene targets of NDD-associated emVars

To experimentally identify target genes of emVars within their endogenous genomic context, we used CROP-seq coupled with CRISPR interference (CRISPRi) to perturb emVar loci. We performed CROP-seq in a hPSC-derived neuron model expressing doxycycline-inducible NGN2 and safe-harbor-integrated dCas9-KRAB. We designed guide RNAs (gRNAs) targeting the 38 emVars identified by MPRA (**Figure 1**, **Methods**). The resulting gRNA library was cloned into a modified CROP-seq vector, packaged into lentivirus, delivered to hPSCs, enriched by fluorescence-activated cell sorting (FACS), and differentiated into neurons as previously described^14^ (**Figure 3A**, **Figure S4A**). During gRNA design, rs148926303 and rs202851 were excluded because efficient targeting gRNAs could not be identified. Following CROP-seq, rs12908208 was further excluded because no cells carrying its targeting gRNAs were detected, whereas rs726024 and rs726025 were treated as a single target locus because they are separated by only 9 bp. The final CROP-seq screen therefore targeted 35 emVars across 34 distinct loci (**Table S4**).

**Figure 3.**
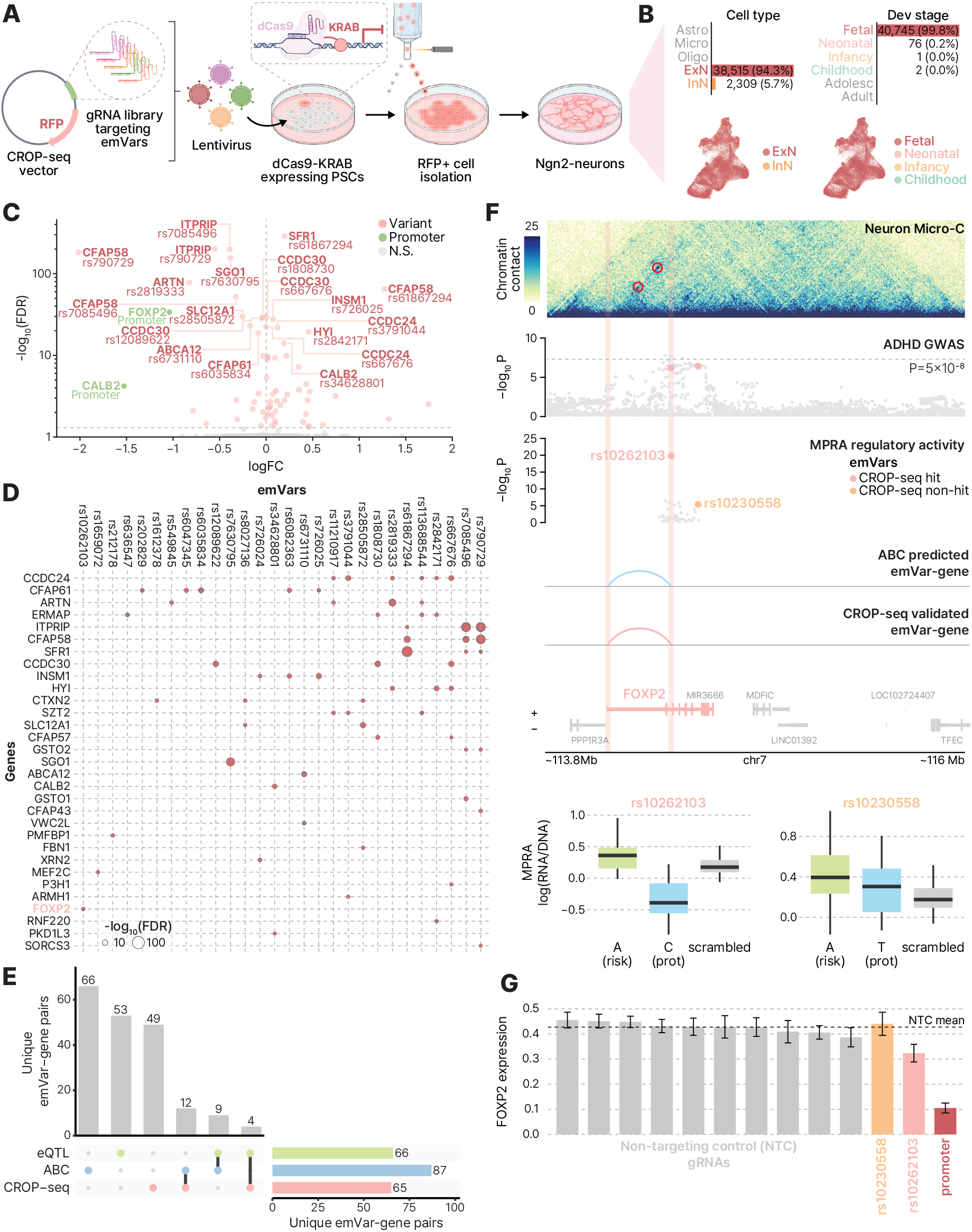
CROP-seq targeting of emVars in hPSC-derived neurons. **A.** Schematic of the CROP-seq workflow. gRNAs targeting emVar loci were cloned into a CROP-seq vector, packaged into lentivirus, delivered to dCas9-KRAB-expressing hPSCs, enriched for red-fluorescent-protein-positive (RFP+) cells by FACS, and differentiated into NGN2 neurons. **B**. CROP-seq targeted hPSC-derived neurons predominantly map to excitatory neurons and the fetal developmental stage. Bar plots show the number and percentage of cells assigned to each major cell type and developmental stage. Astro, astrocytes; Micro, microglia; Oligo, oligodendrocytes; ExN, excitatory neurons; InN, inhibitory neurons. **C**. Volcano plot showing differential expression results from CROP-seq analyses using mZINB. Labeled points indicate significant gene-emVar pairs (pink) and promoter-targeting positive controls (green). **D**. Dot plot of significant CROP-seq-supported gene-emVar pairs. Dot size represents the significance of perturbation effects, with genes shown in rows and emVars shown in columns. **E**. UpSet plot showing overlap among gene-emVar pairs nominated by eQTL mapping, ABC predictions, and CROP-seq. Bars indicate the number of unique gene-emVar pairs in each set or intersection. **F**. *FOXP2* locus plot with normalized Micro-C contact profiles for neurons, ADHD GWAS association statistics, MPRA regulatory activity, ABC-predicted emVar-gene links, CROP-seq-supported emVar-gene links, and gene annotations. Boxplots below show MPRA reporter activity for the risk and protective alleles of rs10262103 (left) and rs10230558 (right) compared with scrambled controls. **G**. *FOXP2* expression following CRISPRi targeting with NTC gRNAs and representative rs10230558-targeting, rs10262103-targeting, and promoter-targeting gRNAs. Bars represent mean-normalized *FOXP2* expression among cells carrying individual gRNAs, with error bars indicating standard error of the mean. The dashed line indicates mean *FOXP2* expression across cells carrying NTC gRNAs.

We next characterized the identity and developmental state of the hPSC-derived neurons using scRNA-seq. Because *NGN2* induction can generate peripheral nervous system (PNS)-like neurons in addition to central nervous system (CNS)-like excitatory neurons^55^, we filtered the dataset to retain high-confidence CNS-like excitatory neurons (**Figure S4B-C, Methods**). To further assess cellular identity, we compared the filtered cells with a human brain single-cell atlas^20^. Among the filtered cells, 94.3% mapped to excitatory neurons and 99.8% mapped to the fetal developmental stage (**Figure 3B**). Together with the developmental chromatin context of emVars (**Figure 1G**, **Figure S2B**) and the modest neuronal bias observed for predicted target genes (**Figure 2D**), these results support the use of hPSC-derived neurons as a relevant neurodevelopmental context for evaluating emVar function at their endogenous loci.

We then assessed the quality of the CROP-seq dataset using the filtered cells. Transcriptomic complexity and standard QC metrics were consistent across sequencing inlets (**Figure S4D-H**). Most neurons were assigned a single gRNA (30,789 cells, 84.6%; **Figure S4I**), with a median of 50 total gRNA unique molecular identifiers (UMIs) per neuron (**Figure S4J**). In addition, 4,210 multiplets meeting predefined compatibility criteria were retained, resulting in 34,999 cells used for downstream CROP-seq analyses (**Figure S4K**, **Methods**). Together, these metrics demonstrated robust gRNA capture and representation across the analyzed neuronal population.

Because expression of many candidate target genes was sparse in the CROP-seq dataset, we compared a standard negative binomial (NB) regression model with a marginalized zero-inflated negative binomial (mZINB) model^56^, which jointly accommodates variation in gene expression and excess zero counts (**Figure S5A-B**). The initial analysis comprised 3,283 candidate gRNA-gene pairs across 350 candidate target genes. Despite substantial numbers of cells per perturbation condition, candidate genes were frequently undetected at the single-cell level: across tests, the median fraction of nonzero expression was 36.7% among perturbed cells and 36.0% among control cells (**Figure S5C**). This sparsity motivated us to evaluate whether explicitly modeling excess zeros could improve detection of perturbation-associated expression changes.

NB and mZINB were applied to the same initial candidate test set and subjected to shared QC criteria (**Figure S5D**). Following the initial filtering, 1,128 tests remained, with subsequent model-specific filtering yielding 792 evaluable tests for mZINB and 825 for NB (**Figure S5D**). After per-gRNA Benjamini-Hochberg correction, mZINB identified 65 significant emVar-gene pairs, compared with 7 identified by NB. Five pairs were shared between the two models, whereas 60 were detected only by mZINB and 2 only by NB (**Figure S5E**). Despite this difference in sensitivity, estimated Wald coefficients were highly concordant between models (Spearman ρ = 0.95; **Figure S5F**), indicating that the models generally agreed on the direction and magnitude of perturbation effects.

The increased sensitivity of mZINB was more apparent for sparsely expressed genes. Significant pairs identified by mZINB involved genes with higher zero fractions than those identified by NB (**Figure S5G**). Moreover, mZINB achieved a higher fitted log-likelihood than NB for 82.3% of evaluated tests (**Figure S5H**). Together, these analyses indicate that accounting for excess zeros improves sensitivity to perturbation effects among sparsely expressed candidate genes while retaining effect-size estimates that are highly concordant with NB.

As positive controls for CRISPRi activity, we also examined gRNAs targeting *FOXP2*, *CALB2*, and *MEF2C* promoters. Cells carrying the promoter-targeting gRNAs showed reduced expression of each target gene relative to non-targeting control (NTC) cells (**Figure S5I**), and all three perturbations were significant under the mZINB likelihood ratio test (**Figure S5J**). In contrast, NB identified significant effects only for *FOXP2* (**Figure S5J**), further illustrating the increased sensitivity of mZINB for detecting perturbation-associated expression changes. Together, these results confirmed effective CRISPRi-mediated repression and support mZINB as a well-suited framework for the sparse, zero-inflated expression structure of the target genes in our CROP-seq dataset. We therefore selected mZINB as the primary model for subsequent CROP-seq analyses.

The 65 CROP-seq-supported emVar-gene pairs involved 31 unique genes (**Figure 3C**, **Table S5**), several of which have established roles in neurodevelopmental and neuronal processes. *INSM1*, for example, encodes a zinc-finger transcription factor with a panneurogenic role in the developing nervous system and promotes the generation of basal progenitors in the developing neocortex^57,58^. *CALB2* encodes the calcium-binding protein calretinin, which is expressed in distinct developing neuronal populations and regulates intracellular calcium dynamics. Notably, reduced densities of calretinin-positive interneurons have been reported in the caudate nucleus of individuals with ASD^59,60^. *ITPRIP* encodes a membrane-associated protein that binds to inositol 1,4,5-trisphosphate receptors (IP3Rs), which mediate intracellular calcium release^61^. Interestingly, reduced IP3R-mediated calcium release has been reported in patient-derived cells from multiple monogenic ASD syndromes, including fragile X syndrome and *TSC1*- and *TSC2*-associated tuberous sclerosis, further linking this signaling pathway to ASD-associated biology^62^. These findings show that endogenous emVar perturbations affect genes involved in diverse neurodevelopmentally relevant processes, including neurogenesis and calcium signaling.

CROP-seq also revealed that emVar-gene relationships were not uniformly one-to-one (**Figure 3D**). While most CROP-seq target genes were affected by a single emVar, a subset responded to perturbation of multiple emVars, with as many as 6 emVars converging on the same gene (**Figure S6A**). Conversely, although most emVars affected a single gene, several perturbations affected multiple genes, with up to 6 target genes identified for an individual emVar (**Figure S6A**). These results indicate that endogenous emVar perturbation can capture both convergent regulation, in which multiple regulatory elements influence a shared target gene, and pleiotropic regulation, in which a single regulatory locus influences multiple genes.

We next quantified the extent to which CROP-seq supported emVar-gene relationships nominated by predictive mapping approaches. Among 144 unique emVar-gene pairs nominated by eQTL or ABC, 16 were also supported by CROP-seq (11.1%), corresponding to 13 unique genes (**Figure 3E, Figure S6B**). These overlapping genes spanned a broad range of baseline expression in our hPSC-derived neurons, with regulatory effects detected even for relatively lowly expressed genes (**Figure S6C**). For example, *HYI* and *ARTN* were each nominated by multiple predictive mapping resources despite relatively low baseline expression, indicating that cross-method support was not restricted to highly expressed genes. Support was more frequent among ABC predictions (12/87, 13.8%) than eQTL predictions (4/66, 6.1%) (**Figure 3E**). Conversely, most CROP-seq-supported pairs were not nominated by either eQTL or ABC (49/65, 75.4%), suggesting that endogenous perturbation can capture regulatory relationships beyond those identified by current variant-to-gene prediction approaches.

The *FOXP2* locus illustrates how CROP-seq can refine candidate regulatory links following MPRA-based variant prioritization and ABC-based gene mapping. This locus contained two emVars, rs10262103 and rs10230558. Although both looped to the *FOXP2* promoter in neuronal chromatin interaction profiles, only rs10262103 was predicted to regulate *FOXP2* via the ABC model (**Figure 2B**). CRISPRi targeting of rs10230558 produced *FOXP2* expression levels indistinguishable from NTC gRNAs, whereas targeting rs10262103 significantly reduced *FOXP2* expression, albeit to a lesser extent than promoter-targeting positive control gRNAs (**Figure 3G**). Consistent with these observations, rs10262103 showed stronger allelic regulatory activity in MPRA and was the only emVar annotated as an enhancer in hPSC-derived neurons, whereas rs10230558 was annotated as quiescent (**Figure 3F-G**; **Figure S2A**). This example highlights the value of combining MPRA, variant-gene mapping, and endogenous perturbation to prioritize regulatory variants and their target gene

### NDD emVar perturbations converge on developmentally regulated transcriptional programs

We next explored whether emVar perturbations converge on broader transcriptional programs. While individual emVars have modest effects on specific target genes, their perturbations could propagate through coordinated gene networks and manifest at the level of transcriptional programs. We reasoned that network-based analyses can capture coordinated perturbation effects that are not readily detected in gene-level analyses, and therefore applied consensus non-negative matrix factorization (cNMF) to gene expression profiles from perturbed cells^63^ (**Figure 4A**, **Methods**). This analysis identified 30 transcriptional programs representing diverse biological processes in hPSC-derived neurons (**Table S6**).

**Figure 4.**
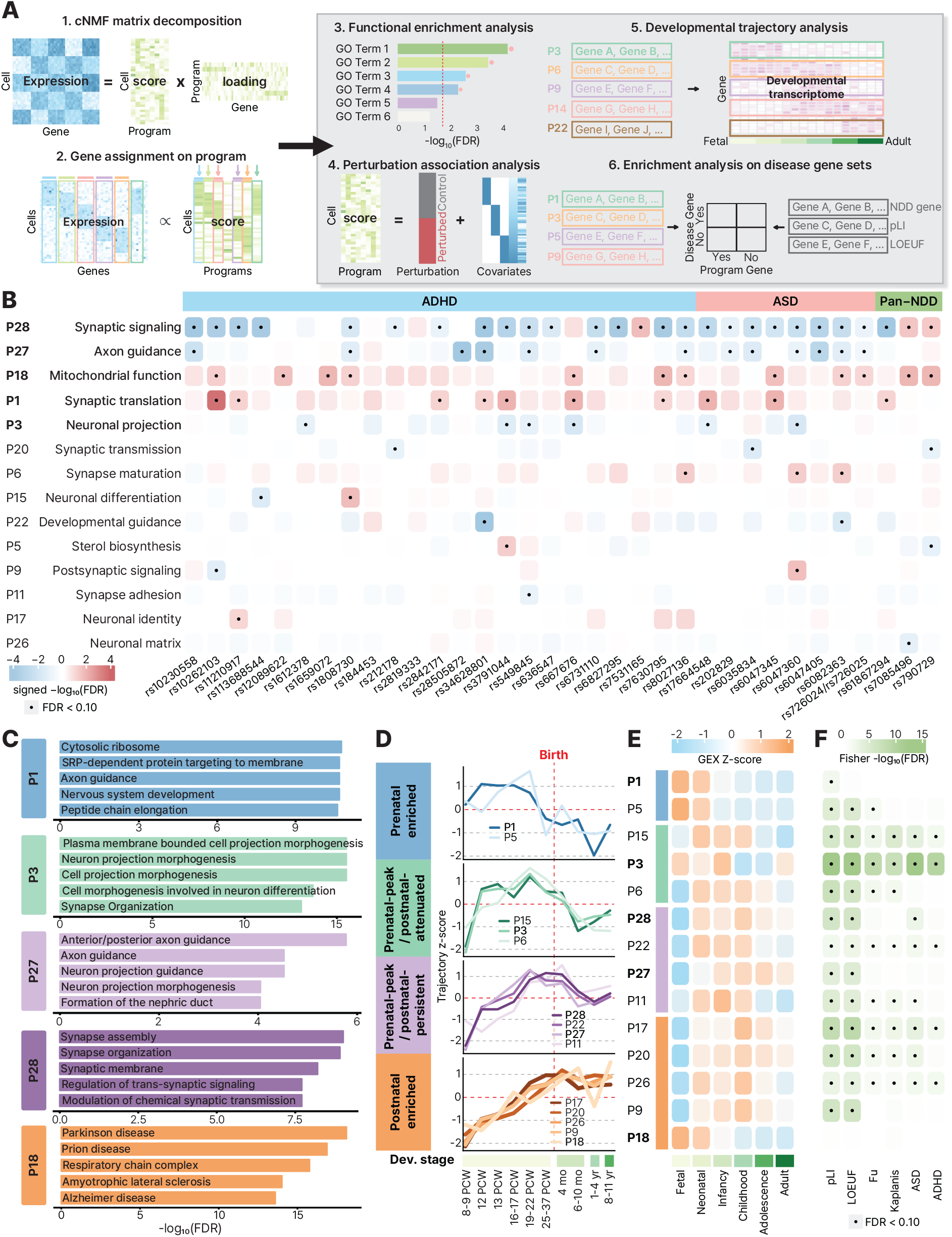
NDD emVar perturbations converge on developmentally regulated neuronal programs. **A.** Schematic of the cNMF-based program analysis workflow. cNMF was applied to gene expression profiles from perturbed cells to decompose the expression matrix into program scores and gene loadings. Genes were assigned to programs, and programs were characterized by functional enrichment, emVar perturbation associations, developmental expression trajectories, and enrichment for disease-associated gene sets. **B**. Heatmap of emVar-associated shifts in program activity upon perturbation of each emVar. Rows indicate emVar-associated programs and their assigned biological annotations. Columns indicate emVars grouped by associated GWAS category. Color represents signed –log_10_(FDR), with blue indicating negative (decreased) program activity and red indicating positive (increased) program activity upon emVar perturbation. Dots indicate associations passing FDR < 0.1. Convergent programs are shown in bold. **C**. Top functional enrichment terms for convergent programs. Bars indicate the significance of the top 5 enriched terms used to assign biological annotations to convergent programs. **D.** Mean cortical developmental trajectories of emVar-associated programs. Programs are grouped according to their temporal expression pattern: prenatal enriched, prenatal-peak/postnatal-attenuated, prenatal-peak/postnatal-persistent, and postnatal enriched. The dashed red line indicates birth. Dev. stage, developmental stage; PCW, postconception weeks; mo, months; yr, years. **E**. Excitatory neuronal developmental trajectories of emVar-associated programs. Values indicate within-program gene-expression z-scores across six excitatory neuronal developmental stages. **F**. Enrichment of emVar-associated programs for LoF-intolerant genes (pLI ≥ 0.9 and LOEUF < 0.35), ASD risk genes (Fu), DD risk genes (Kaplanis), and ASD/ADHD risk genes (NDDGeneHub). Color represents enrichment significance, and dots indicate enrichment passing FDR < 0.1.

Of these, we retained 14 programs that showed at least one significant emVar-associated perturbation shift (**Figure 4B**). These programs spanned 34 targeted emVar loci and comprised 85 significant emVar-program associations (**Table S7**), of which 53 were negative (i.e., emVar perturbation was associated with decreased program activity). Among the 14 retained programs, we identified 5 convergent programs (P1, P3, P18, P27, P28), defined as programs significantly regulated by ≥ 6 emVars at FDR < 0.1 (**Figure 4B**). The strongest convergence was observed for P28 and P27, which together accounted for 40 of 85 significant associations (47.1%), including 37 negative associations. In contrast, P18 and P1 accounted for 22 associations (25.9%), all of which were positive, while P3 contained 6 significant associations, all of which were negative (**Figure 4B**).

We performed pathway analysis to assign biological identities to the convergent programs. To annotate each program, we prioritized dominant themes across Gene Ontology (GO), KEGG, and Reactome, then used Synaptic Gene Ontology (SynGO) to refine annotations related to neuronal and synaptic function when relevant^64–66^ (**Figure 4C**, **Figure S7A-B**, **Table S8**). P28 was annotated as a synaptic signaling program based on enrichment for synaptic organization and trans-synaptic signaling. P27 was annotated as an axon guidance program with enrichment for axon guidance and neuron projection guidance pathways. P1 was enriched for translation at synaptic compartments and was annotated as a synaptic translation program. P3 was enriched for neuronal projection morphogenesis as well as synaptic structure and vesicle regulation, consistent with a neuronal projection program. P18 was enriched for mitochondrial respiratory-chain and oxidative-phosphorylation processes and was therefore annotated as a mitochondrial function program. P18 was also enriched for KEGG pathways associated with multiple neurodegenerative diseases, consistent with emerging evidence suggesting that individuals with ASD exhibit signatures of accelerated aging^67,68^ (**Figure 4C**, **Table S8**).

We next asked whether these emVar-associated programs showed coordinated developmental trajectories. To this end, we leveraged bulk developmental brain RNA-seq^69^ data to reconstruct mean cortical expression trajectories for each program spanning early fetal development through childhood (**Figure 4D**). The emVar-associated 14 programs separated into 4 distinct temporal patterns. P1 and P5 showed prenatal enrichment, peaking before birth and declining sharply around the time of birth. P3, P6, and P15 were enriched during fetal development and attenuated after the neonatal period, exhibiting a prenatal-peak/postnatal-attenuated pattern. P11, P22, P27, and P28 peaked prenatally but remained expressed into later development, exhibiting a prenatal-peak/postnatal-persistent pattern. In contrast, P9, P17, P18, P20, and P26 showed postnatal enrichment. These trajectories indicate that emVar-associated programs are not only biologically distinct, but also temporally structured across brain development, suggesting that NDD risk variants may perturb pathways that are preferentially active during different windows of neurodevelopmental vulnerability.

We next examined whether these developmental patterns were preserved at cellular resolution using a single-cell transcriptomic atlas of the developing human cortex^20^ (**Figure 4E**). Because emVar-associated programs were defined from hPSC-derived neurons, we focused on their developmental trajectories within the excitatory neuronal lineage. The resulting trajectories broadly recapitulated the temporal organization observed in the bulk brain data. Programs assigned to earlier developmental patterns, including P1, P5, and P3, generally showed higher relative expression during fetal, neonatal, or infant stages, whereas programs assigned to later developmental patterns showed relatively greater expression during postnatal stages. Programs with prenatal-peak/postnatal-persistent trajectories, including P11, P22, P27, and P28, maintained expression across a broader developmental window. Although the precise timing of peak expression differed slightly between the bulk and single-cell datasets for some programs, the overall early-versus-late developmental organization was largely preserved.

Examination across broader neural and glial lineages further showed that most programs were most prominently expressed within neuronal lineages, particularly excitatory and inhibitory neurons, while several exhibited more restricted expression in non-neuronal populations (primarily oligodendrocytes) at specific developmental stages (**Figure S7C**). P1 and P18 showed comparatively broader early developmental expression across multiple lineages. Together, these analyses indicate that emVar-associated transcriptional programs exhibit reproducible developmental organization across bulk and single-cell datasets while retaining distinct lineage-specific expression patterns.

One important question is whether gene programs perturbed by common variants converge on genes implicated by rare coding mutations, suggesting shared biological pathways through which common and rare genetic variants contribute to NDD risk. We therefore tested whether emVar-associated programs were enriched for genes that harbor rare variants associated with ASD, ADHD, and developmental disorders (DD)^1,70–72^. Multiple emVar-associated programs showed significant enrichment for genes implicated by rare coding variation (**Figure 4F**, **Table S9**). Among the convergent programs, P3 exhibited the strongest enrichment, overlapping ASD^1,70–72^, ADHD^70,73^, and DD^1,70–72^ risk genes. Representative genes contributing to this enrichment included *ARID1B*, *AUTS2*, *MBD5*, and *MYT1L*. P28 also showed enrichment for ASD risk genes^70,73^, including *BCL11A*, *CASZ1*, *CNTNAP2*, *NRXN1*, *PLCB1*, and *RPS6KA3* (**Figure 4F**).

We also examined whether emVar-associated programs are likely to be dysregulated in postmortem brains of ASD-affected individuals^74^. Out of the convergent programs, P3, P27, and P28 were enriched for differentially expressed genes (DEGs) in excitatory neurons from ASD postmortem brains (**Figure S7D**), suggesting that common variants may perturb neurodevelopmental programs that leave lasting transcriptional signatures in the adult ASD brain.

Previous studies have shown that genes implicated in NDDs are frequently dosage sensitive and exhibit intolerance to disruptive mutations. We therefore further evaluated enrichment of loss-of-function (LoF) intolerant genes using pLI^75^ (probability of being LoF intolerant) and LOEUF^71^ (loss-of-function observed/expected upper bound fraction) metrics. All convergent programs except P18 were enriched for LoF intolerant genes.

Collectively, these findings suggest that NDD emVars converge on dosage-sensitive, developmentally regulated neuronal programs that are enriched for genes affected by rare coding mutations and dysregulated in ASD brains, linking common genetic risk to shared molecular mechanisms underlying NDDs.

## DISCUSSION

Here, we systematically interrogated the regulatory consequences of common genetic variants previously associated with NDDs by integrating two complementary high-throughput functional approaches. MPRA enabled scalable screening of GWAS variants and prioritized 38 emVars with regulatory activity. We then mapped these variants to candidate target genes using eQTL- and ABC model-based approaches. We subsequently used CROP-seq to perturb nearly the entire set of MPRA-prioritized emVars in hPSC-derived neurons. This integrated strategy connected genetic association to variant regulatory activity, predicted target genes, and endogenous transcriptional consequences within a neurodevelopmentally relevant cellular context.

Several broader principles emerged from integrating these functional layers. First, NDD emVars showed strong developmental context dependence, including greater overlap with fetal brain chromatin states, enrichment of Alu elements, and disruption of TFBMs linked to neuronal differentiation and signaling. Second, integration of MPRA and CROP-seq provides a framework for moving from scalable variant prioritization to endogenous target gene discovery. MPRA enabled scalable prioritization of variants with allele-dependent regulatory activity, whereas CROP-seq tested their transcriptional consequences within the endogenous genomic context. Third, endogenous perturbation supported only a subset of predicted emVar-gene relationships captured by either eQTL or ABC, highlighting the incompleteness of current variant-gene mapping strategies. Finally, emVar perturbations extended beyond cis target genes to broader transcriptional programs involving synaptic signaling, axon guidance, neuronal projection, and mitochondrial function. These programs followed distinct developmental trajectories and were enriched for dosage-sensitive genes, rare NDD-associated genes, and genes dysregulated in ASD brains. Together, these findings suggest that while common NDD-associated regulatory variants may act through heterogeneous proximal targets, they converge on shared, developmentally regulated molecular programs.

The TFBMs altered by NDD emVars highlighted upstream regulatory pathways that may shape their developmental effects. Nominally enriched motifs implicated transcriptional regulators with established roles in neural progenitor maintenance^24^ (SOX family), neuronal differentiation^76,77^ (SIN3A), nuclear receptor signaling^25,27,78,79^ (RARA, RARG, RORA), and mitochondrial regulation (ERR family). Importantly, several biological themes implicated by TFBM analysis re-emerged among the downstream consequences of endogenous emVar perturbation. Retinoic acid signaling regulates neural differentiation, neuronal patterning, and axonal outgrowth, paralleling the enrichment of P27 and P3 for axon guidance and neuronal projection-related processes^27,78^. Similarly, emVars altered motifs for estrogen-related receptors (ERR family) that regulate mitochondrial and metabolic programs^28,80^, while P18 was independently enriched for respiratory-chain and oxidative-phosphorylation genes. The RFX family provides another convergent axis: RFX2 and RFX3 regulate ciliogenesis and neural development^81,82^, and CROP-seq identified multiple cilia-associated genes among downstream emVar targets. Our independent CROP-seq study of schizophrenia-associated genes also identified ciliary transcriptional programs and linked their perturbations to altered primary cilia morphology^83^. Together, these findings suggest that upstream regulatory signals disrupted by NDD emVars can converge with downstream transcriptional consequences on shared neurodevelopmental pathways.

Finally, the improved performance of mZINB motivated us to generalize this modeling strategy into ZiPert (**Z**ero-**I**nflated Modeling for **Pert**urbation Differential Expression Analysis), a framework that supports multiple count models for identifying perturbation-associated expression changes in sparse single-cell data, including standard NB and zero-inflated models (https://github.com/thewonlab/ZiPert). ZiPert provides a flexible analytical tool for single-cell perturbation datasets, allowing investigators to apply mZINB to lowly expressed genes, while retaining standard NB models for genes with less sparse expression. In parallel, we linked cNMF-derived transcriptional programs to emVar perturbation status, providing a complementary approach for assessing broader trans-regulatory consequences beyond individual target genes. Together, these analytical strategies enable interrogation of perturbation effects across both gene-level and program-level scales in sparse single-cell datasets.

## LIMITATIONS

Several limitations should be considered when interpreting these findings. MPRA was performed in NPCs, whereas CROP-seq was conducted in hPSC-derived neurons, and regulatory effects may differ across developmental stages and cell types. In addition, episomal MPRA measures regulatory activity outside the native chromosomal context, whereas CRISPRi perturbs endogenous regulatory elements but does not reproduce the precise allele-specific sequence change, so the two assays capture complementary but distinct aspects of variant function. Current variant-gene mapping approaches also remain incomplete, as eQTL and ABC predictions differ in developmental context and regulatory assumptions, and only a subset of predicted relationships were supported by CROP-seq, while many CROP-seq-supported relationships were not nominated by either approach. Furthermore, our CROP-seq analysis focused primarily on genes whose nearer start or end boundary lay within 500 kb of the targeted site; more distal or indirect effects may therefore have been missed. Finally, this study primarily characterized transcriptomic phenotypes and did not directly assess downstream effects on neuronal morphology, physiology, excitability, or network function. Future studies incorporating matched developmental contexts, allele-specific genome editing, improved cell type-specific variant-gene mapping, and higher-order neuronal phenotypes will be important for extending these findings.

## AUTHOR CONTRIBUTIONS

H.W., A.G., and J.C.M. designed the study. J.C.M. constructed MPRA libraries and conducted MPRA experiments with assistance from J.L.B. J.C.M. also led MPRA data analysis with assistance from H.L. (data preprocessing), Y.G. (gene mapping), and S.L. (TFBM analysis). T.U.A. generated Micro-C libraries from hiPSC-derived NPCs and neurons under the supervision of Z.F. and A.S.B. E.H. processed and analyzed Micro-C data. N.Y. and R.S. generated hPSCs expressing dCas9-KRAB and *NGN2*. A.G. performed CROP-seq experiments with assistance from S.J. H.M. developed the ZiPert package, and A.G. and H.M. jointly analyzed CROP-seq data. A.G., J.C.M., H.M., and H.W. generated figures. A.G., J.C.M., H.M., and H.W. co-wrote the manuscript, which was subsequently reviewed and edited by the rest of the authors.

## ACKNOWLEDGEMENTS

We thank members of the Won lab, as well as Dr. John S. Preisser, for helpful discussions and comments about this paper. This research was supported by the PsychENCODE consortium (R01MH122509, H.W.), the Impact of Genomic Variation on Function (IGVF) consortium (UM1HG012003, H.W., A.S.B.), the National Institute of General Medical Sciences (5T32GM067553, S.L; 5T32GM135128, J.C.M., A.G.), the Genomics of ASD: Pathways to Genetic Therapies award (1012863, H.W., N.Y.) and the Data Analysis Award (SFI-AN-AR-Data Analysis-00019464-01, H.W.) from the Simons Foundation Autism Research Initiative (SFARI).

## RESOURCE AVAILABILITY

### Lead contact

Any inquiries about the project, analytical results, or other information should be directed to Hyejung Won.

### Data and code availability

Raw and processed MPRA data, CROP-seq data, and micro-C data have been deposited in the NIMH Data Archive (NDA) and are publicly accessible under NDA Collection ID C6209 (Collection DOI: 10.15154/113x-dx35). Access to these data is governed by the NDA Data Use Certification and can be requested through the NDA portal. Custom codes for MPRA analysis (crossdisorder-MPRA) and (cross_disorder_MPRA_count_matrix_pipeline), activity-by-contact (ABC)-based variant-gene prediction pipeline (ACV2G), and CROP-seq differential expression analysis (ZiPert) is available on our lab GitHub page.

## DECLARATION OF INTERESTS

The authors declare no competing interests.

## STAR METHODS

### EXPERIMENTAL MODEL DETAILS

#### Primary human NPC line used for MPRA experiments

MPRA experiments were performed using primary NPCs derived from donor #88, a genetically male donor at gestational week 14. The acquisition, derivation, and culture of these NPCs has been previously described^13,14^. NPCs were maintained under neural progenitor growth conditions prior to MPRA transfection as described below.

#### hiPSC line used for Micro-C experiments

Micro-C experiments were performed using the UNC001-A hiPSC line, which was generated from peripheral blood of a healthy female donor using integration-free reprogramming. The generation and characterization of UNC001-A has been previously described^84^. UNC001-A hiPSCs were differentiated into neural progenitor cells and cortical neurons as described below, and three isogenic biological replicates of each differentiated cell type were used for Micro-C library generation.

#### hPSC line used for CROP-seq experiments

CROP-seq experiments were performed using the dCas9-KRAB H1 human embryonic stem cell line, derived from the genetically male H1 hESC line obtained from WiCell. The line was engineered to constitutively express dCas9-KRAB from the AAVS1 safe-harbor locus. An AAVS1-targeting donor vector carrying an FRT-flanked EMCV-IRES-neomycin selection cassette and EF1α-driven dCas9-HA-KRAB was co-transfected with a plasmid expressing Cas9 and an AAVS1-targeting gRNA (Addgene #80494). Following targeted integration, transient Flp recombinase expression was used to remove the neomycin selection cassette. This engineered line has been previously described^85^. Doxycycline-inducible NGN2 was subsequently introduced using the piggyBac Tet-ON vector PB-TO-oNGN2 (Addgene, Cat. No. #198397).

## METHOD DETAILS

### MPRA library construction

The MPRA library construction was carried out as previously described^14^. Briefly, the Agilent oligonucleotide library was diluted to 2 pM and PCR-amplified using MPRA-chip-primer-F and MPRA-chip-primer-R with Q5 Hot Start High-Fidelity 2X Master Mix (NEB, Cat. No. M0494). The amplified library was purified using the DNA Clean & Concentrator-5 kit (Zymo Research, Cat. No. D4004). Twenty-base-pair barcodes were subsequently introduced by PCR using MPRA-bc-primer-F and MPRA-bc-primer-R with NEBNext 2X Q5 HiFi Hot Start Master Mix, followed by an additional purification step using the DNA Clean & Concentrator-5 kit.

The barcoded oligonucleotide library and the MPRA vector Donor_eGP2AP_RC (Addgene plasmid #133784) were digested with SpeI-HF (NEB, Cat. No. R3133S) and MluI-HF (NEB, Cat. No. R3198S). The digested plasmid was isolated from a 1% agarose gel using the Zymoclean Gel DNA Recovery Kit (Zymo Research, Cat. No. D4008), and the digested oligonucleotide library was ligated into the linearized vector using T7 DNA Ligase (NEB, Cat. No. M0318S). The ligation mixture was immediately transformed in six separate reactions of Endura Electrocompetent Cells (Lucigen, Cat. No. 60242-2) and recovered in the supplied recovery medium for 1 h at 37 °C. Transformed bacteria were plated on 10-cm LB agar plates containing kanamycin and incubated overnight at 37°C. Colonies were collected and expanded overnight in 4 L of LB containing kanamycin at 30°C, after which plasmid DNA was purified using the ZymoPURE II Plasmid Maxiprep Kit (Zymo Research, Cat. No. D4203).

To establish variant-barcode associations, variant sequence-barcode fragments were amplified from the plasmid library using BC_Map_P5_R and BC_Map_P7_F_Ind_# with NEBNext 2X Q5 HiFi Hot Start Master Mix, thereby adding P5 and P7 Illumina adapters and sample indices. PCR products were purified and size-selected using SparQ beads (VWR, Cat. No. 76302-834) and sequenced on an Illumina NovaSeq SP platform using 2 × 150-bp reads, 20% PhiX, and the custom sequencing primers BC_Map_R1seq, BC_Map_R2seq, and BC_Map_Index. Barcodes were assigned to their corresponding variant-containing sequences using custom code as previously described (see Code Availability)^14^.

After barcode mapping, the minimal promoter (minP)-luciferase fragment was excised from pMPRAdonor2 (Addgene plasmid #49353) using KpnI-HF (NEB, Cat. No. R3142S) and XbaI (NEB, Cat. No. R0145S) and gel-purified using the Zymoclean Gel DNA Recovery Kit (Zymo Research, Cat. No. D4008). The barcoded Donor_eGP2AP_RC library was digested with the same enzymes, purified, and ligated to the minimal promoter-luciferase fragment using T7 DNA Ligase (NEB, Cat. No. M0318S). The ligation mixture was transformed into Endura Electrocompetent Cells (Lucigen, Cat. No. 60242-2), and transformants were selected and expanded in kanamycin-containing medium as described above. The final MPRA plasmid library was purified using the ZymoPURE II Plasmid Maxiprep Kit (Zymo Research, Cat. No. D4203).

### MPRA library transfection into human NPCs

The sourcing, generation, culture, and MPRA transfection of the human NPCs used in this study have been previously described^14^. Briefly, genetically male NPCs obtained at gestational week 14 (donor #88) were seeded at 8 × 10^5^ cells per well in 6-well plates coated with poly-L-ornithine (10 μg/mL; Sigma-Aldrich, Cat. No. P3655-100MG) and laminin (5 μg/mL; Gibco, Cat. No. 23017015). NPCs were maintained in Neurobasal A medium (Thermo Fisher Scientific, Cat. No. 10888022) supplemented with Primocin (100 μg/mL; InvivoGen, Cat. No. ant-pm-2), BIT 9500 (10%; STEMCELL Technologies, Cat. No. 09500), GlutaMAX (1%; Thermo Fisher Scientific, Cat. No. 35050061), heparin (1 μg/mL; Sigma-Aldrich, Cat. No. H3393-100KU), fibroblast growth factor (FGF; 20 μg/mL; Thermo Fisher Scientific, Cat. No. 100-18B), LIF (2 ng/mL; PeproTech, Cat. No. 300-05), EGF (20 μg/mL; PeproTech, Cat. No. AF-100-15), and PDGF (20 ng/mL; PeproTech, Cat. No. 100-00AB).

Forty-eight hours after seeding, cells received a half-medium change using the same medium containing twice the standard concentration of growth factors and nucleic acid transfection enhancer (NATE; 1:50 final dilution; InvivoGen, Cat. No. lyec-nate). Each well was then transfected with 2.5 μg of the MPRA plasmid library and 1 μg of pUC19 plasmid DNA (Thermo Fisher Scientific, Cat. No. SD0061) using Lipofectamine STEM (Invitrogen, Cat. No. STEM00015) according to the manufacturer’s instructions. Cells received another half-medium change 48 h after transfection. At 72 h after transfection, DNA and RNA were isolated using the Quick-DNA/RNA Miniprep Plus Kit (Zymo Research, Cat. No. D7003). For each well, cells were lysed in 600 μL DNA/RNA Shield and 600 μL lysis buffer.

### MPRA DNA and RNA library preparation

Following nucleic acid extraction, cDNA was synthesized from each RNA sample using SuperScript IV Reverse Transcriptase (Invitrogen, Cat. No. 18090050) with the custom reverse-transcription primer Lib_Hand_RT, which anneals downstream of the barcode sequence. The resulting cDNA was PCR-amplified using Lib_Hand and Lib_seq_Luc_R with Q5 Hot Start High-Fidelity 2X Master Mix (NEB, Cat. No. M0494), and the amplified product was purified using the DNA Clean & Concentrator-25 kit (Zymo Research, Cat. No. D4033).

In parallel, an aliquot of each DNA sample was PCR-amplified using Lib_Hand and Lib_seq_Luc_R with NEBNext 2X Q5 HiFi Hot Start Master Mix and purified using the DNA Clean & Concentrator-25 kit. P5 and P7 Illumina adapter sequences and sample indices were then added to both cDNA- and DNA-derived libraries by PCR using P5_seq_Luc_F and P7_Ind_#_Han. Following amplification, libraries were purified and size-selected using AMPure XP beads (Beckman Coulter, Cat. No. A63881) with 0.7X and 0.9X bead ratios. Final libraries were sequenced on an Illumina NovaSeq SP platform using a custom 35 × 8 × 0 × 0 (R1 × I7 × I5 × R2) cycle configuration, 20% PhiX, and R1_seq and Index_seq custom sequencing primers.

### Reprogramming of peripheral blood cells for Micro-C

Peripheral blood mononuclear cells (PBMCs) were isolated from 10 mL of whole blood collected in 3.2% sodium citrate. Erythroblasts were expanded from PBMCs for 14 days and reprogrammed using integration-free Sendai virus vectors encoding OCT4/POU5F1, SOX2, KLF4, and c-MYC, as previously described^84^. Cells were subsequently cultured on inactivated mouse embryonic fibroblasts (MEFs). Colonies displaying typical pluripotent stem cell morphology emerged approximately 2 weeks after transduction and were manually selected and expanded on feeders for three passages. Cells were then transferred to Matrigel-coated culture dishes (Corning, Cat. No. 354277) and transitioned to StemFlex medium (Thermo Fisher Scientific, Cat. No. A3349401). Established iPSC cultures were maintained on Matrigel with daily medium replacement and passaged every 3–5 days using a 50mM EDTA (Corning, Cat. No. 46-034-CI) dissociation buffer in PBS (Gibco, Cat. No. 14190-144).

### Cortical neuron differentiation for Micro-C

iPSCs were differentiated into cortical neurons using an adapted dual-SMAD inhibition protocol^86^. Undifferentiated iPSCs were dissociated with Accutase (Sigma, Cat. No. A6964), and 3 × 10^5^ cells were cultured in suspension in StemScale PSC medium (Thermo Fisher Scientific, Cat. No. A4965001) supplemented with 10 µM Y-27632 (Selleckchem, Cat. No. S1049) on an orbital shaker at 37°C and 5% CO2. After 48 h, cell aggregates were dissociated with Accutase, and 3 × 10^5^ cells were replated in suspension for neural induction in StemScale PSC medium supplemented with 10 µM Y-27632, 1.5 µM CHIR99021 (Selleckchem, Cat. No. S1263), 20 µM SB431542 (Selleckchem, Cat. No. S1067), and 50 nM LDN-193189 (Selleckchem, Cat. No. S2618). CHIR99021 was withdrawn after 24 h, and cultures were maintained under neural induction conditions for 10 days with daily medium replacement. Neurospheres were dissociated approximately twice weekly at a 1:3 ratio.

Neural progenitor cells were subsequently expanded for 7 days in neuronal expansion medium consisting of Advanced DMEM/F12 (Thermo Fisher Scientific, Cat. No. 12634010) supplemented with Neural Induction Supplement (Thermo Fisher Scientific, Cat. No. A1647801) and 20 ng/mL FGF (Thermo Fisher Scientific, Cat. No. 100-18B). For terminal neuronal differentiation, NPCs were plated onto poly-D-lysine- (Thermo Fisher Scientific, Cat. No. A3890401) and laminin-coated (Sigma, Cat. No. CC095) culture surfaces and maintained in neuronal maturation medium composed of Neurobasal (Thermo Fisher Scientific, Cat. No. 21103049) supplemented with GlutaMAX (Thermo Fisher Scientific, Cat. No. 35050061), B27 supplement (Thermo Fisher Scientific, Cat. No. 17504001), N2 supplement (Thermo Fisher Scientific, Cat. No. 17502001), MEM non-essential amino acids (Thermo Fisher Scientific, Cat. No. 11140050), β-mercaptoethanol (Thermo Fisher Scientific, Cat. No. 21985-023), insulin (Sigma, Cat. No. I9278), 10 ng/mL brain-derived neurotrophic factor (BDNF; Thermo Fisher Scientific, Cat. No. 450-02), 10 ng/mL glial cell line-derived neurotrophic factor (GDNF; Thermo Fisher Scientific, Cat. No. 450-10), and 10 µM DAPT (Selleckchem, Cat. No. S2215). Neuronal cultures were maintained for 45 days after differentiation.

### Micro-C library generation

Micro-C libraries were generated from iPSC-derived neural progenitor cells and cortical neurons, with three isogenic biological replicates per cell type. Libraries were prepared using the Dovetail Micro-C kit (Dovetail Genomics, Cat. No. PN 20101E). Cells were sequentially cross-linked with 3 mM disuccinimidyl glutarate (DSG; Thermo Fisher Scientific, Cat. No. A35392) and 1% formaldehyde diluted in PBS from 16% formaldehyde stock (Thermo Fisher Scientific, Cat. No. 28906), followed by micrococcal nuclease digestion and SDS-mediated lysis. Chromatin fragments were captured on Chromatin Capture Beads, end-repaired, biotinylated, and proximity-ligated. Following reversal of cross-links and DNA purification, Illumina paired-end sequencing adapters were added. Biotinylated ligation products were enriched using streptavidin beads, PCR-amplified, and size-selected prior to sequencing. Libraries were sequenced on an Illumina NovaSeq 6000 using 2 × 150-bp paired-end reads, generating approximately 1.8 billion read pairs.

### CROP-seq target selection and gRNA design

Following identification of the 38 MPRA emVars, we attempted to design two 20-bp gRNAs targeting each emVar locus (Table S4). Candidate gRNAs were designed within ±150 bp of the variant position using Benchling. Guide specificity and potential off-target activity were subsequently evaluated using BLAT^87^ and GuideScan2^88^. For these QC analyses, the 20-bp protospacer and corresponding protospacer-adjacent motif (PAM) were evaluated together as a 23-bp sequence. Candidate gRNAs were excluded if BLAT identified an identical 23-bp match at another genomic location or if GuideScan2 returned a “multiple perfect matches found” error. emVars for which no gRNA satisfying these criteria could be identified were excluded from the CROP-seq screen. For guides retained for cloning, the PAM sequence was removed and the 20-bp protospacer was flanked by the 5′ sequence TGGAAAGGACGAAACACCG and the 3′ sequence GTTTTAGAGCTAGAAATAGCAAGTTAAAATAAGGC for compatibility with the CROP-seq-tRFP cloning workflow.

For positive controls, three CRISPRi gRNAs targeting the transcription start site of each of FOXP2, CALB2, and MEF2C were selected using CHOPCHOP^89^. Negative controls consisted of one GFP-targeting gRNA and nine non-targeting control gRNAs selected from a previously established CRISPR screening control set^90^.

### CROP-seq library cloning

Guide oligonucleotides were synthesized by Integrated DNA Technologies (IDT) as 74-nt single-stranded DNA oligonucleotides containing the 19-bp 5′ cloning sequence, 20-bp gRNA protospacer, and 35-bp 3′ cloning sequence described above. Oligonucleotides were individually resuspended and normalized prior to pooling.

The CROPseq-tRFP vector (Addgene, Cat. No. #230938) was digested with BsmBI-v2 (NEB, Cat. No. R0739S) in NEBuffer r3.1 at 55°C for 1 h, followed by heat inactivation at 80°C for 20 min. Digested vector was subsequently treated with recombinant shrimp alkaline phosphatase (rSAP; NEB, Cat. No. M0371S) at 37°C for 1 h and heat-inactivated at 65°C for 5 min. The reaction was resolved on a 0.8% agarose gel, and the approximately 8.3-kb vector backbone was excised and purified using the Zymoclean Gel DNA Recovery Kit (Zymo Research, Cat. No. D4008).

For Gibson assembly, individual gRNA oligonucleotides were pooled at equal abundance and diluted to a final pooled concentration of 100 nM. Each assembly reaction contained 11 fmol of digested CROPseq-tRFP backbone, 200 fmol of pooled gRNA oligonucleotides, and NEBuilder HiFi DNA Assembly Master Mix (NEB, Cat. No. E2621S). Reactions were incubated at 55°C for 1 h and purified using the DNA Clean & Concentrator-5 Kit (Zymo Research, Cat. No. D4004).

Purified assembly products were electroporated into Endura Electrocompetent Cells (Lucigen, Cat. No. 60242-2), with up to 50 ng of assembled plasmid introduced per 25 μL of competent cells. Following electroporation, cells were recovered for 1 h at 37°C with shaking at 225 rpm and plated on LB agar containing ampicillin. Serial dilutions were plated to estimate library representation, and libraries achieving >200 bacterial colonies per gRNA were expanded for plasmid preparation. Colonies were pooled and expanded in LB containing ampicillin, and plasmid DNA was isolated by maxiprep using the QIAfilter Plasmid Maxi Kit (Qiagen, Cat. No. 12263).

### CROP-seq library diversity QC

Following plasmid library amplification, cloning quality was initially assessed by restriction digestion with MluI-HF (NEB, Cat. No. R3198S) and electrophoresis on a 0.8% agarose gel. Libraries were subsequently evaluated by sequencing to assess gRNA representation. The gRNA-containing region was amplified from 10 ng of plasmid library using LibQC_i5 and indexed LibQC_i7 primers with NEBNext Q5 Hot Start HiFi PCR Master Mix (NEB, Cat. No. M0543L) and SYBR Green (Invitrogen, Cat. No. S7563). The optimal PCR cycle number was determined by qPCR as the cycle at which fluorescence reached one-quarter of the amplification plateau. Four preparative PCR reactions were then performed using the empirically determined cycle number, pooled, and purified using the DNA Clean & Concentrator-5 Kit (Zymo Research, Cat. No. D4004). Amplicon size was assessed by TapeStation D1000 (Agilent, Cat. No. 5067-5582 and 5067-5583), with an expected product of approximately 300 bp. Libraries were sequenced on an Illumina MiSeq Nano with a 40% PhiX spike-in, and gRNA abundance was quantified to evaluate representation of individual guides prior to lentiviral packaging. Libraries passing representation QC were submitted to the UNC Lenti-shRNA Core Facility for lentiviral production.

### Lentiviral delivery of CROP-seq library to hPSCs

The pooled CROP-seq lentiviral library was delivered to dCas9-Krab/H1 hPSCs when the cells were at approximately 50% confluency. The lentiviral preparation had a functional titer of 1.44 × 10^10^ transducing units (TU)/mL and was diluted 1:10 in DMEM/F12 (Thermo Fisher Scientific, Cat. No. 11330032) prior to addition to the cells. Cells were then transduced at a multiplicity of infection (MOI) of 1. After 24 h, the lentivirus was removed by complete media replacement. Cells were subsequently expanded prior to fluorescence-activated cell sorting for tRFP-positive cells. An untreated well was maintained in parallel as a negative control for establishing the tRFP sorting gate.

Once cultures reached approximately 80–90% confluency, cells were washed with PBS (Corning, Cat. No. 21-040-CV) and dissociated with Accutase (Thermo Fisher Scientific, Cat. No. A1110501) at 37°C for 4–5 min. Cells were gently triturated to generate a single-cell suspension, diluted in StemFlex medium, and pelleted at 200 × g for 5 min. Cell pellets were resuspended in StemFlex supplemented with 10 μM Y-27632 ROCK inhibitor and cryopreserved in a StemFlex-based freezing medium containing 10% DMSO (Sigma, Cat. No. D4540). Cryopreserved cells were subsequently thawed on the day of FACS.

### Dissociation and FACS of CROP-seq-transduced hPSCs

Following lentiviral transduction and expansion, hPSCs were enriched for CROP-seq library integration by FACS based on tRFP expression. Cells were cryopreserved prior to sorting and then thawed in StemFlex medium (Thermo Fisher Scientific, Cat. No. A3349401) and pelleted at 200 × g for 5 min on the date of the sort. Cells were resuspended in StemFlex pre-sort medium supplemented with 2% fetal bovine serum (FBS; Corning, Cat. No. 35-015-CV), 1% Antibiotic-Antimycotic (Thermo Fisher Scientific, Cat. No. 15240062), and 10 μM Y-27632 dihydrochloride ROCK inhibitor (Tocris, Cat. No. 1254). Cell suspensions were then passed through a 40-μm cell strainer (Corning, Cat. No. 352340) to remove debris and aggregates. DNase I from the Worthington Papain Dissociation System (Worthington, Cat. No. LK003153) was added to a final concentration of 5 μg/mL together with 5 mM MgCl₂ (Thermo Fisher Scientific, Cat. No. AM9530G) to minimize cell aggregation during sorting. For live-cell discrimination, cells were stained with Calcein AM (Thermo Fisher Scientific, Cat. No. C3099) at a final concentration of 100 nM and incubated for 15–20 min prior to sorting.

Lentivirus-transduced cells and untreated control cells were transported on ice to the UNC Flow Cytometry Core for sorting. Untreated hPSCs were used to establish the tRFP-negative gate, and live, Calcein-positive, tRFP-positive cells were isolated by FACS. Sorted cells were collected into StemFlex post-sort medium supplemented with 4% FBS, 2% Antibiotic-Antimycotic, and 20 μM Y-27632 dihydrochloride ROCK inhibitor. Following sorting, cells were transported on ice and plated with post-sort medium onto Matrigel-coated 24-well plates using Corning Matrigel Growth Factor Reduced Basement Membrane Matrix (Corning, Cat. No. 354230) for recovery and subsequent expansion prior to neuronal differentiation. A half-medium change with StemFlex and 1% Antibiotic-Antimycotic was performed 48 h after sorting. Cultures showing continued recovery and expansion received a complete medium change the following day, whereas slower-recovering cultures were maintained for an additional 24 h before initiation of daily complete medium changes. Antibiotic-Antimycotic supplementation was maintained for 7 days after sorting and replating, after which cells were cultured with StemFlex alone and subsequent medium changes were performed without Antibiotic-Antimycotic.

### Neuron differentiation of CROP-seq-transduced hPSCs

CROP-seq-transduced dCas9-Krab/H1 hPSCs were differentiated into glutamatergic neurons using an NGN2-based differentiation protocol that was previously described^14,91^. Following recovery and expansion after tRFP-positive cell sorting, hPSCs were dissociated with Accutase (Thermo Fisher Scientific, Cat. No. A1110501) and seeded on Matrigel-coated 6-well plates at 7.5 × 10^5^ cells per well on day −3.

For pre-differentiation, cells were maintained in KnockOut DMEM/F12 (Thermo Fisher Scientific, Cat. No. 12660012) supplemented with 1× MEM non-essential amino acids (Thermo Fisher Scientific, Cat. No. 11140050), 1× N2 supplement (Thermo Fisher Scientific, Cat. No. 17502048), 10 ng/mL NT-3 (PeproTech, Cat. No. 450-03), 10 ng/mL BDNF (Thermo Fisher Scientific, Cat. No. 450-02), 1 μg/mL laminin (Thermo Fisher Scientific, Cat. No. 23017015), and 2 μg/mL doxycycline (Sigma, Cat. No. D3447). Y-27632 ROCK inhibitor was added at 10 μM during day −3 plating only and omitted from subsequent pre-differentiation medium changes. Half-medium changes with pre-differentiation medium were performed on days −2 and −1, and cells were maintained under these conditions until day 0.

On day 0, cells were dissociated with Accutase, pelleted at 200 × g for 5 min, and resuspended in N2/B27 differentiation medium consisting of a 1:1 mixture of DMEM/F12 with HEPES (Thermo Fisher Scientific, Cat. No. 11330032) and Neurobasal-A (Thermo Fisher Scientific, Cat. No. 10888022), supplemented with 1× MEM non-essential amino acids, 0.5× GlutaMAX (Thermo Fisher Scientific, Cat. No. 35050061), 0.5× N2 supplement, 0.5× B27 minus vitamin A (Thermo Fisher Scientific, Cat. No. 12587010), 10 ng/mL NT-3, 10 ng/mL BDNF, 1 μg/mL laminin, and 2 μg/mL doxycycline. Y-27632 ROCK inhibitor was added at 10 μM during day 0 plating only. Cells were seeded onto poly-D-lysine-coated plates (Thermo Fisher Scientific, Cat. No. A3890401) at approximately 1 × 10^6^ cells per well of a 6-well plate and maintained under neuronal differentiation conditions through day 13. On day 13, neurons were collected for single-cell dissociation and FACS.

#### Dissociation and FACS of CROP-seq-transduced hPSC-derived Neurons

On day 13 of neuronal differentiation, CROP-seq-transduced hPSC-derived neurons were dissociated for FACS using a papain-based dissociation protocol^91^. Conditioned neuronal medium was collected from each well for neuron resuspension prior to sorting and centrifuged at 200 × g for 5 min to pellet dead cells. Papain and DNase I from the Worthington Papain Dissociation System (Worthington, Cat. No. LK003153) were prepared separately according to the manufacturer’s kit components. Papain was resuspended in 1× Hanks’ Balanced Salt Solution (Thermo Fisher Scientific, Cat. No. 24020117) to a final concentration of 20 U/mL and pre-warmed at 37°C for 10 min. DNase I was resuspended in 1× HBSS at 1 mg/mL and added immediately before use to a final concentration of 5 μg/mL, together with 5 mM MgCl_2_ (Thermo Fisher Scientific, Cat. No. AM9530G). Neuronal cultures were washed with PBS (Corning, Cat. No. 21-040-CV), and 500 μL of dissociation solution (composed of papain, DNase, and MgCl_2_) was added per well of a 6-well plate. Cells were incubated at 37°C for 10 min, after which the dissociation reaction was quenched with five volumes of DMEM/F12 with HEPES (Thermo Fisher Scientific, Cat. No. 11330032) supplemented with 10% FBS. Neuronal sheets were gently triturated to generate a single-cell suspension.

Cells were pelleted at 200 × g for 10 min and resuspended in 500 μL conditioned neuronal medium supplemented with 2% FBS (Corning, Cat. No. 35-015-CV) and 10 μM Y-27632 dihydrochloride ROCK inhibitor (Tocris, Cat. No. 125410). Cell suspensions were passed through a 40-μm cell strainer (Thermo Fisher Scientific, Cat. No. 087711) and stained with Calcein AM (Thermo Fisher Scientific, Cat. No. C3099) at a final concentration of 100 nM for 15–20 min for live-cell discrimination. Lentivirus-transduced neurons and RFP-negative control cells were transported on ice for FACS. Live, Calcein-positive, tRFP-positive neurons were isolated and collected into conditioned neuronal medium supplemented with 4% FBS and 20 μM Y-27632 ROCK inhibitor for subsequent single-cell processing.

### Single-cell gene expression and gRNA enrichment library preparation

Single-cell cDNA was generated by the UNC Advanced Analytics Core from three CROP-seq inlets across two experimental batches using the 10x Genomics Chromium Single Cell 3′ v4 workflow, with one inlet processed in the first batch and two inlets processed in the second batch. For the first experimental batch, the corresponding gene-expression library was also prepared by the UNC Advanced Analytics Core. For the second experimental batch, single-cell cDNA from the two inlets was returned for subsequent gene-expression library preparation prior to sequencing.

Following generation of the single-cell cDNA, gRNA-derived sequences were separately enriched using a two-step PCR strategy with custom GuideEnrich primers. For PCR1, five 20-μL reactions were prepared per inlet using 50 ng of cDNA per reaction, NEBNext Q5 Hot Start HiFi PCR Master Mix (NEB, Cat. No. M0543), and 0.5 μM each of GuideEnrich_1 and GuideEnrich_2. One reaction was supplemented with SYBR Green and used for qPCR to determine the optimal amplification cycle number at one-quarter of the fluorescence plateau. The remaining four reactions were amplified using the empirically determined cycle number, with PCR1 limited to a maximum of 10 cycles to minimize overamplification. All three inlets were amplified for 10 cycles. PCR1 products were pooled and purified using the DNA Clean & Concentrator-5 Kit (Zymo Research, Cat. No. D4004).

For PCR2, the entire purified PCR1 product from each inlet was used as a template with 0.5 μM each of indexed GuideEnrich_3 and GuideEnrich_4 primers and NEBNext Q5 Hot Start HiFi PCR Master Mix. A separate qPCR reaction was again used to determine the amplification cycle number at one-quarter of the fluorescence plateau. A minimum of eight PCR2 cycles was used to ensure complete addition of sequencing adapters; all three inlets were amplified for eight cycles. PCR1 and PCR2 used an initial denaturation at 98°C for 30 s, followed by cycles of 98°C for 10 s, 69°C for 20 s, and 65°C for 15 s, with a final extension at 65°C for 2 min. PCR2 products were size-selected using sparQ PureMag Beads (Quantabio, Cat. No. 95196) with sequential 0.5× and 0.7× bead ratios to enrich fragments approximately 300–700 bp in length. Final gRNA-enrichment libraries were quantified using the Qubit 1X dsDNA High Sensitivity Assay Kit (Thermo Fisher Scientific, Cat. No. Q33231) and assessed by TapeStation prior to sequencing.

### Sequencing Pooled Gene Expression and gRNA Enrichment Libraries

For the first experimental batch, the gene-expression library prepared by the UNC Advanced Analytics Core and the corresponding gRNA enrichment library were combined at an 80:20 molar ratio, respectively, and sequenced through the UNC High-Throughput Sequencing Facility (HTSF) using a NovaSeq 1.5B full flow cell.

For the second experimental batch, gene expression libraries from the two CROP-seq inlets were pooled equimolarly, and their corresponding gRNA-enrichment libraries were pooled separately at equimolar abundance. The resulting gene expression and gRNA enrichment pools were then combined at an 80:20 molar ratio, respectively, and sequenced on one lane of an Illumina NovaSeq X Plus 25B flow cell.

### QUANTIFICATION AND STATISTICAL ANALYSIS

#### Variant Selection for MPRA Testing

Variants were selected from 18 genome-wide significant loci identified across three neurodevelopmental disorder GWAS datasets: ASD^8,9^, ADHD^10^, and pan-NDD^12^. Within each locus, variants with GWAS association P < 1 × 10^-5^ were considered for MPRA testing, as previously described^14^. Variants containing recognition sites for restriction enzymes used during downstream library cloning (MluI, SpeI, KpnI, or XbaI) were excluded. This yielded 729 variant loci prior to MPRA mapping quality control. For each selected variant, a 150-bp genomic sequence centered on the variant was extracted from the hg19 reference genome and synthesized as part of the MPRA oligonucleotide library. The library additionally included 6 positive-control sequences consisting of 3 tiled fragments from each of the CMV and EF1α promoters, as well as 200 scrambled negative-control sequences GC-matched to the variant library. Following barcode-mapping quality control, loci with a mapping-error proportion ≥ 0.30 were excluded, leaving 727 variant loci for downstream MPRA analysis.

#### MPRA data preprocessing

MPRA sequencing data were generated and processed as part of the larger MPRA dataset previously described^14^. Briefly, barcode DNA libraries were sequenced to an average depth of 81 million reads, and barcode RNA libraries were sequenced to an average depth of 167 million reads. Barcode-variant associations established during barcode-mapping sequencing were used to generate barcode reference files. Variant-barcode reads were then aligned using BWA-MEM (version 0.7.17-r1188)^92^, and RNA and DNA count matrices were generated using featureCounts (version 2.0.6)^93^ with --minOverlap = 20, --fracOverlap = 1, and --fracOverlapFeature = 1. Barcodes mapping to more than one variant were excluded. RNA and DNA counts were then aggregated across barcodes for each reporter sequence.

Barcode-level quality control was performed as previously described^14^. Outlier barcodes were defined as those with log2(RNA/DNA) values greater than the median log_2_(RNA/DNA) for the corresponding variant plus 2 and were excluded. Variants represented by fewer than five barcodes per allele or lacking measurements for both risk and protective alleles were also excluded. Finally, variants were removed if, for either allele, more than 30% of total DNA and RNA reads originated from barcodes potentially attributable to another sequence when mismatches were permitted. The present study subsequently restricted the processed MPRA dataset to the NDD-associated loci described above, yielding 729 loci prior to mapping-quality filtering and 727 loci retained for downstream MPRA analyses.

#### MPRA Reproducibility Analysis

To assess reproducibility across the 22 biological replicates, RNA and DNA counts were summed across barcodes for each MPRA reporter construct within each replicate. Reporter activity was summarized as log[(RNA + 1)/(DNA + 1)]. Pairwise Pearson correlation coefficients were calculated between these log-ratio profiles across all 22 replicates, yielding 231 pairwise comparisons. Positive controls, scrambled negative controls, and variant-containing reporters were retained in the count matrix and annotated by reporter type. For visualization of MPRA activity, sample-level log ratios were averaged across the 22 replicates for each of the 1,454 allele-specific reporters and the control reporters.

#### Defining MPRA-active Elements and EmVars

MPRA element activity and allelic effects were quantified using the mpra package (version 1.24.0)^94^ as previously described^14^. Allelic effects were tested at the variant-locus level. Allelic P values were adjusted for multiple testing using Benjamini-Hochberg (BH) across 729 loci prior to mapping-quality filtering. Loci with a mapping-error proportion ≥ 0.30 were excluded from reported calls, leaving 727 loci for downstream analysis. Variants were classified as allelic if they had an allelic FDR < 0.05 and a mapping-error proportion < 0.30.

Element activity was evaluated separately for both allele-specific reporter sequences at each locus, yielding 1,454 reporter tests across the 727 retained loci. Reporter activity was assessed using limma::treat (version 3.56)^95^ with a nonzero effect-size threshold of log_2_(1.238016) ≈ 0.308. Reporter-level P values were adjusted together across all 1,454 tests using the BH procedure. A locus was classified as MPRA-active if at least one of its two allele-specific reporters showed a positive activity logFC and reporter FDR < 0.10. Thus, locus-level activity was defined as the union of the two allele-specific reporter calls rather than by selecting the most significant allele. Finally, emVars were defined using both element activity and allelic regulatory effects. Thirty loci were both MPRA-active and significantly allelic. An additional eight loci that were significantly allelic but did not meet the MPRA-active criterion were retained as emVars because they exhibited strong allelic effects, defined as |allelic logFC| > 0.75. These two mutually exclusive groups yielded a final set of 38 emVars, which were used for downstream analysis.

#### Defining Background SNPs

To generate a matched genomic background for characterization of NDD emVars, we first defined the set of MPRA-tested NDD variants and their associated LD regions. Candidate background SNPs were required to fall outside the tested MPRA variants and their associated LD regions and to show no significant GWAS association (P > 0.05). Background SNPs were matched to tested NDD variants by minor allele frequency and LD, with both metrics required to fall within ±10% of the corresponding value for the tested variant. For each tested variant, up to 10 matched background SNPs were randomly sampled when more than 10 eligible SNPs were available. Matched SNPs were then combined across variants and duplicate entries were removed. This procedure yielded a fixed background set of 6,145 matched SNPs, originally represented in the hg19 genome assembly.

#### Chromatin Accessibility of emVars Across Cortical Cell Types and Developmental Stages

We characterized chromatin accessibility of the 38 finalized emVars using publicly available human cortical single-cell ATAC-seq data^20^. Accessibility signal was summarized in 150-bp windows centered on each emVar. Finalized accessibility matrices were generated for four broad cortical cell-type groups (astrocytes, excitatory neurons, inhibitory neurons, and oligodendrocytes) and six excitatory-neuron developmental stages (fetal, neonatal, infancy, childhood, adolescence, and adulthood).

For emVar accessibility heatmaps, pheatmap (version 1.0.12)108 was used. Accessibility values were standardized within each emVar across the cell types or developmental stages. Variants were ordered by hierarchical clustering using a distance metric of 1 − Pearson correlation and complete-linkage clustering. Z-score clipping used in the heatmaps was applied for visualization only.

#### ChromHMM Annotation and Enrichment Analysis

To characterize the chromatin states associated with NDD emVars, the 38 emVars and 6,145 matched background SNPs were analyzed using 15-state ChromHMM annotations^17^. Variants and background SNPs were represented in hg19 coordinates and intersected with ten brain and neural ChromHMM tracks for annotation. Eight of these contexts were included in the emVar–background comparison. Individual Roadmap states were consolidated into seven broader functional categories as follows: Enh and EnhG were classified as enhancer; Tx, TxWk, and TxFlnk as transcribed; ReprPC, ReprPCWk, and Het as repressive; TssA and TssAFlnk as active promoter; TssBiv, BivFlnk, and EnhBiv as poised/bivalent; ZNF/Rpts as ZNF/repeats; and Quies as quiescent.

For each chromatin-state category within each of the eight neural or brain contexts, the proportion of emVars assigned to that state was compared with the proportion of matched background SNPs using two-sided Fisher’s exact tests. Multiple-testing correction was performed using the BH procedure across all 56 state-by-context comparisons.

#### Repeat Element Overlap Analysis

To characterize overlap of NDD emVars with repetitive genomic elements, 150-bp windows centered on each hg19 variant (±75 bp) were intersected with hg19 RepeatMasker annotations ^96^. Repeat elements were grouped into five major classes: Alu, SVA, L1, LTR, and hAT-Charlie elements, using repeat-name patterns defined in the analysis pipeline. A variant was considered to overlap a given repeat class when more than 50% of its 150-bp window overlapped an annotation belonging to that class.

For each repeat class, the proportion of the 38 emVars overlapping that class was compared with the corresponding proportion among the 6,145 matched background SNPs using a two-sided Fisher’s exact test. P values were adjusted for multiple testing using the BH procedure across the five repeat-class comparisons.

#### Transcription Factor Binding Motif Analysis

We assessed predicted allele-dependent TF-binding motif effects for the 38 finalized emVars and a motif-specific set of 377 GWAS-nonsignificant SNPs matched on minor allele frequency and linkage disequilibrium. Candidate background SNPs were restricted to variants with GWAS P > 0.05 located outside ±500 kb of the emVars. Up to 10 background SNPs per emVar were sampled within the same chromosome using minor allele frequency and LD matching tolerances of 10%, 20%, 50%, and 100%. Sampling used a fixed random seed of 20260416 and yielded 377 unique background SNPs. Using human MotifDb (version 1.50.0) ^97^ motifs excluding the stamlab source, we ran motifbreakR (version 2.22.0)^21^ on hg19 alleles with filterp = TRUE, an information-content method, a threshold of 1 × 10^-4^, and equal nucleotide background frequencies. We retained motif calls classified by motifbreakR as effect = “strong”, harmonized TF aliases and accession-based names to canonical symbols, and counted each TF–variant pair once across motif providers.

For each TF, enrichment of qualifying motif effects among emVars relative to matched background SNPs was assessed using a one-sided Fisher’s exact test. The 33 nominally enriched TF motifs (unadjusted P < 0.05) were displayed; none remained significant after BH correction. For visualization, odds ratios were calculated after adding a pseudocount of 1 to each contingency-table cell. Fisher’s exact test P values were calculated from the original counts without pseudocounts. For the supplementary motif gain/loss analysis, motif direction was oriented relative to the GWAS risk allele using allele-specific motif scores, with motif creation defined when the risk allele had a higher motif score than the comparison allele and disruption defined when the risk allele had a lower score.

#### Transcription Factor Expression Context Across Neural Differentiation

Expression of transcription factors represented in the motif analysis was assessed using the LIBD Stem Cell Browser^54^. Raw gene-level counts were used to calculate full-transcriptome library sizes for each sample, and expression was converted to log(CPM + 1) before selecting the transcription factors of interest. The displayed comparison included NPC samples (n = 20) and the NEURONS_PLUS_ASTROS condition (n = 31), labeled as “Neuron” in the figure. TF symbols were standardized prior to matching with the expression matrix, and duplicate expression rows mapping to the same TF symbol were averaged within the sample. All 33 nominally enriched TFs were successfully matched to the expression dataset.

For each TF, mean log(CPM + 1) expression was calculated across samples within each condition, with standard errors derived from the sample-level values. For the scaled-expression heatmap, mean expression was first calculated across all six differentiation conditions in the LIBD dataset and then standardized within each TF across conditions. Only NPC and NEURONS_PLUS_ASTROS values were displayed in the final heatmap.

#### emVar-Gene Mapping via eQTL

We integrated the 38 finalized emVars with three brain eQTL resources representing distinct developmental and cellular contexts: adult brain MetaBrain eQTLs^35^, fetal brain eQTLs^34^, and adult single-nucleus brain (SingleBrain) eQTLs^36^. Variant-gene associations were matched to emVars using rsID and genomic position where available, with analyses harmonized to GRCh38 coordinates. Candidate gene identities were tracked using stable Ensembl gene identifiers, with gene symbols retained for display.

For the fetal brain dataset, associations with FDR < 0.05 were retained. For SingleBrain, associations with Random_FDR < 0.05 were retained. Adult MetaBrain associations were prefiltered at FDR < 0.05, and no additional significance filter was applied during integration with the finalized emVar set. To compare regulatory direction between MPRA and eQTL datasets, eQTL effect alleles were aligned to the MPRA allele encoding. Effect-size signs were reversed where necessary so that all eQTL coefficients were oriented to MPRA A2. Directionally concordant associations were defined as those in which the aligned eQTL effect and MPRA allelic log fold change had the same sign, and only these concordant associations were retained for downstream emVar–gene nomination.

#### Micro-C Data Processing

Micro-C reads were processed using the 4D Nucleome (4DN) Hi-C processing pipeline^98^. Sequencing reads were aligned to the GRCh38 reference genome using BWA^92^. Ligation pairs were identified using pairtools parse and sorted using pairtools sort^99^. Sequencing replicates were merged within each biological replicate according to the 4DN workflow, and PCR duplicates were removed using pairtools dedup. Deduplicated interaction pairs were binned into 10-kb contact matrices using Cooler^100^. Contact matrices were visualized using plotgardener^101^.

#### Variant-Gene Mapping via the ABC model

We used an activity-by-contact-based approach to prioritize variant-gene relationships separately in NPCs and cortical neurons (CNs). For each MPRA variant, protein-coding genes with GENCODE v33 promoters located within ±1 Mb were considered candidate targets. Chromatin accessibility at each variant was quantified using cell-type-matched ATAC-seq BigWig tracks: the GSE291907 day-6 NPC ATAC-seq track^38^ for NPCs and a pooled NeuN-positive neuronal ATAC-seq track^37^ for CNs. Accessibility was summarized over a 500-bp window centered on each variant. Chromatin contact was obtained from the corresponding NPC or CN Micro-C data at 10-kb resolution, using the contact frequency between the variant-containing bin and promoter-associated bins. When multiple promoter bins or contact records supported the same variant-gene pair, the maximum contact count was used.

For each variant-gene pair, both the ATAC-seq accessibility signal and Micro-C contact frequency were log1p-transformed, and the transformed values were multiplied to generate an ABC score. Scores were standardized separately for each gene across the corresponding GWAS candidate variants to generate gene-specific ABC z-scores. High-confidence activity-contact mappings were defined as variant-gene pairs with finite ABC z-scores in the upper 10% of the overall z-score distribution within each cell-type dataset. These analyses were performed using Activity-Contact Variant-to-Gene Prioritization (ACV2G), an R-based computational pipeline developed for this study and publicly available at https://github.com/thewonlab/ACV2G. The resulting ABC predictions were subsequently restricted to the 38 finalized emVars for integration with the eQTL-based emVar-gene mappings.

#### Expression Context of emVar-Nominated Target Genes

Expression of emVar-nominated target genes across neural differentiation was assessed using previously published transcriptomic data from the LIBD Stem Cell Browser^54^. Library-size normalization was performed using the complete transcriptome prior to subsetting to the 60 genes nominated by eQTL and/or ABC mapping. Genes were matched to the expression dataset using Ensembl gene identifiers, and normalized expression values were subsequently summarized for the nominated gene set. The condition displayed as “Neuron” corresponds to the NEURONS_PLUS_ASTROS condition in the source dataset.

For NPC-versus-neuron comparisons, mean expression for each nominated gene was summarized as log(CPM + 1). Expression across additional neural differentiation states was standardized gene-wise for visualization in the corresponding heatmap.

#### CROP-seq data pre-processing and cellwise quality control

Sequencing data were processed using 10x Genomics Cell Ranger^102^ (version 9.0.0) with the GRCh38-2024-A reference to generate gene-expression and gRNA UMI count matrices. Cells identified by the standard Cell Ranger cell-calling pipeline were used for Seurat-based preprocessing. For each inlet, genes detected in at least three cells and cells with at least 200 detected genes were retained. Cells with mitochondrial, ribosomal, or MALAT1 transcript fractions exceeding 15%, 20%, or 20%, respectively, were excluded. Cells with total RNA UMI counts exceeding the inlet-specific 99th percentile calculated from guide singlets were also excluded. Among cortical cells passing QC, the distributions of detected genes, RNA UMI counts, mitochondrial and ribosomal transcript fractions, and UMI counts for the most abundant gRNA were examined by inlet using violin plots.

Gene-expression matrices were normalized using Seurat (version 5.3.0)^103^, followed by selection of 2,000 highly variable genes, scaling, and principal component analysis (PCA). Fifty principal components were computed, and batch effects were corrected using Harmony (version 2.0.5)^104^. The first 30 dimensions of the Harmony-corrected embedding were used to generate a two-dimensional UMAP^105^. Cells were clustered at a resolution of 0.8.

#### CROP-seq cell type classification and filtering

Because NGN2-based differentiation can generate PNS-like neuronal populations in addition to CNS-like excitatory neurons, we characterized cell identity within the CROP-seq dataset prior to downstream analysis. Neuronal populations were classified as CNS-like or PNS-like based on their transcriptional state, supported by expression of canonical neuronal and regional marker genes. CNS-like cells showed greater representation of cortical/excitatory-associated markers, including SLC17A6, GRIN2A, and GRIA2, whereas PNS-like cells showed greater representation of peripheral/sensory markers, including ISL1, RET, and PRPH. Progenitor/pluripotency markers (POU5F1, SOX2, NANOG), glial markers (GFAP, AQP4, S100B), and pan-neuronal markers (MAP2, TUBB3, STMN2) were also examined to further characterize the two populations. PNS-like cells were excluded, and the CNS-like population was retained for downstream CROP-seq analyses.

To independently characterize the identity and developmental state of the retained CNS-like neurons, we also performed reference-based annotation using the human prefrontal cortex single-cell atlas^20^. Broad cell-type and developmental-stage identities were assigned using SingleR (version 2.10.0)^106^. Among annotated retained cells, 94.3% mapped to excitatory neurons and 99.8% mapped to the fetal developmental stage, supporting the interpretation of the retained population as predominantly fetal-like excitatory neurons.

#### Assignment of gRNAs to cells

A gRNA was assigned to a cell if at least five UMIs corresponding to that gRNA were detected. Cells with exactly one gRNA meeting this threshold were classified as guide singlets, whereas cells with two or more qualifying gRNAs were classified as guide multiplets. For multiplet classification, assayed variants were grouped into proximity loci by chromosome, with successive variants assigned to the same locus when separated by less than 1 Mb. Multiplets were then categorized according to the targets of the two gRNAs with the highest UMI counts as targeting the same variant, different variants within the same locus, different loci, or positive-control genes. For cis variant-gene analyses, multiplets targeting the same variant or different loci were retained, whereas those targeting different variants within the same locus were excluded. Retained multiplets were included in the perturbation group for each relevant guide based on the complete list of assigned gRNAs. Cells containing guides targeting positive-control genes were excluded from variant analyses. For positive-control analyses, we included only singlets carrying a positive-control gRNA and multiplets containing that gRNA together with an NTC gRNA.

#### Variant-element quality control and cis candidate-gene selection

Before model fitting, guide-to-variant assignments were harmonized across the CROP-seq library. Of the 38 MPRA-defined emVars, rs148926303 and rs202851 were excluded during gRNA design because no suitable targeting gRNAs satisfying the design and specificity criteria could be identified. rs12908208 was excluded because no cells carrying its targeting gRNAs were detected. rs726024 and rs726025 were treated as a single perturbation element because the two variants are separated by only 9 bp and share overlapping guide-targeting windows. Candidate genes were restricted to the same chromosome and selected within 500 kb of the guide target coordinate, with distance measured to the nearer gene start or end boundary. Candidate guide-gene tests were subsequently evaluated based on guide detection and gene-expression properties in perturbation and control cells.

We tested whether expression of these candidate genes differed between emVar-perturbed cells and NTC cells. This restricted cis testing strategy differs from conventional perturbation-based differential expression analyses, which often test approximately 20,000 genes per perturbation and are driven mainly by genes with sufficient expression. Because our candidate set was intentionally limited to nearby genes, it could include low-abundance or cell-state-specific targets for which zero counts were common. We therefore used a mZINB model to explicitly model zero inflation while preserving an interpretable perturbation effect on the covariate-adjusted marginal mean (Supplementary Fig. S5A). Differential expression analysis was performed using ZiPert (https://github.com/thewonlab/ZiPert), an in-house software implementation of mZINB regression for single-cell CRISPR perturbation data.

#### ZiPert model specification

Each guide was compared individually against NTC cells. The perturbation group contained variant singlets and multiplets according to the eligibility criteria described above. The control group was restricted to pure NTC singlets. The full mZINB and NB models included guide-positive status as the main predictor and adjusted for inlet, log(nCount_RNA), log1p(gRNA UMI count), mitochondrial read fraction, S phase score, and G2M phase score. These covariates were included to account for batch/inlet effects, library size, guide capture, mitochondrial content, and cell-cycle state.

#### Model initialization and optimization

Because direct optimization of zero-inflated models can be sensitive to initialization, we used a staged fitting strategy (Supplementary Fig. S5B). For each guide-gene test, we first fit the NB model using the same design matrix to obtain stable initial estimates for the mean-model coefficients and dispersion-related parameters. These NB estimates were then used as warm-start values for the corresponding mZINB fit. The mZINB likelihood was implemented in PyTorch, and model parameters were optimized by directly maximizing the log-likelihood, equivalently minimizing the negative log-likelihood. Optimization was performed with the L-BFGS algorithm, which was used for both the full model containing the guide term and the reduced model without the guide term.

#### Hypothesis testing and effect direction

For each guide-gene test, the mZINB model was compared with a reduced model that removed the guide term but retained the same covariate structure (Supplementary Fig. S5B). The primary significance statistic was a LRT, which evaluates whether including the guide term improves model fit. We used the LRT rather than the Wald p-value for hit calling because the Wald statistic depends directly on an individual coefficient and standard error, which can become unstable for sparse or low-expression genes. After assessing significance with the LRT, the direction of effect was assigned from the sign of the perturbation coefficient estimated in the full mZINB model. This full-versus-reduced comparison is conceptually similar to the model comparison used in hurdle-model frameworks such as MAST, but the mZINB formulation directly accommodates overdispersion and zero inflation while estimating a marginal mean effect.

#### Model filtering and hit calling

We applied sequential QC filters to remove unreliable model fits before hit calling (Supplementary Fig. S5D). Specifically, we retained only successful fits and excluded tests with fewer than five nonzero cells in either perturbation or control cells, negative likelihood ratios, abnormally large Wald standard errors (Wald_SE ≥ 100), missing mapped gene symbols, or non-protein-coding target genes. Multiple testing correction was performed using the Benjamini-Hochberg procedure within each guide, and significant guide-gene effects were defined as those with an LRT-based FDR < 0.05. Variant-gene summaries were generated by collapsing guide-level results according to the variant-element map defined during guide-coordinate QC.

#### Gene Mapping Comparison

Predicted variant–gene pairs were matched to CROP-seq results using rsIDs and stable Ensembl gene IDs. For each mapping resource, support rates were calculated among predicted pairs with an eligible guide-gene test after the QC filters described above. A pair was considered supported if at least one corresponding guide-gene test was significant at FDR < 0.05. Predicted pairs without an eligible test were excluded from the support-rate denominator.

Overlap among eQTL, ABC, and CROP-seq emVar-gene mapping results was visualized with UpSet plots constructed using ggplot2 (version 3.5.2)^107^ and patchwork (version 1.3.0)^108^. At selected loci, plotgardener (version 1.14.0)^101^ was used to display gene-mapping evidence alongside MPRA results, GWAS association statistics, and Micro-C interaction data.

#### Consensus non-negative matrix factorization analysis

Because the direct effect of a variant on individual gene expression can be modest and sparse, we first used cNMF (version 1.6.0) to define recurrent biological gene expression programs in the single-cell transcriptome, and then tested whether each variant altered the usage of these programs. This approach evaluates variant effects at the level of co-regulated gene modules or cell-state programs rather than at the level of individual genes.

cNMF was performed using 3,000 highly variable genes from the normalized expression matrix of CNS-like neurons retained after CROP-seq preprocessing. We evaluated K = 10, 20, 30, 50, and 100 using 200 factorization iterations per K, a random seed of 14, and a consensus density threshold of 0.1. K = 30 was selected based on program stability and reconstruction error. We then used the gene spectra and usage matrix from the K = 30 solution to integrate top-loading genes, GO annotations, and perturbation–program associations. All downstream heatmaps, developmental trajectory analyses, disease gene-set enrichment analyses, and convergence analyses used the K = 30 programs.

#### Assigning genes to cNMF programs

For downstream analyses, we defined a representative gene set for each cNMF program. Although one common strategy is to use a fixed number of top genes from the gene expression program loadings provided by cNMF, assigning the top 300 genes to every program can include genes whose relationship to program usage and actual expression patterns is weak. We therefore directly calculated the genome-wide Spearman correlation between each gene’s expression and the program usage score to define program-specific gene sets.

For each program, genes were ranked by the absolute value of the Spearman correlation. In the main analysis, genes with an absolute Spearman correlation of |ρ| ≥ 0.1 were selected as program genes. Because enrichment results can become unstable when gene sets are too small or too large, programs with fewer than 30 genes passing this threshold used the top 30 genes by absolute correlation as a fallback, including genes below the correlation threshold, whereas programs with more than 100 genes were capped at the top 100 genes by absolute correlation. As a sensitivity check, we also generated gene sets using a |ρ| >= 0.2 threshold, but the main figures and downstream enrichment analyses were centered on the K=30, |ρ| >= 0.1 gene sets. This threshold was selected to balance program specificity and gene set size while maintaining stable downstream enrichment analyses.

#### Functional enrichment analysis

GO enrichment analysis was performed using the gost function from the R package gprofiler2 (version 0.2.4)^109^. Spearman-based gene sets for each program were used as queries, and the default ontology sources included GO biological process (GO:BP), GO molecular function (GO:MF), GO cellular component (GO:CC), KEGG and Reactome (REAC). Analyses used the default annotated-gene background without a custom background gene set. P-values were adjusted using g:Profiler’s default g:SCS method, and terms with adjusted P < 0.05 were considered significant. Terms returned for each program were ranked by p-value. To prevent overly broad biological terms from dominating the annotations, terms with a term size of 800 or more were excluded from figure-level GO plots. Representative annotations for each program were assigned by prioritizing the most strongly enriched significant terms. Enrichment results were summarized as top-term bar plots and program-term heatmaps.

Synaptic Gene Ontology (SynGO; version 1.3)^66^ enrichment was evaluated across all 30 cNMF programs. P29 had no SynGO-annotated query genes and was therefore not tested. Enrichment was evaluated using one-sided Fisher’s exact tests with the SynGO brain-expressed background supplemented with SynGO-annotated genes in the query set. Terms were required to contain at least three overlapping query genes. P-values were adjusted using the BH procedure separately within each program and ontology domain (biological process or cellular component). For the five convergent programs, up to five significant terms with the lowest adjusted P values were displayed per program in Supplementary Figure S7B. All significant SynGO terms across the tested programs are provided in Table S8.

#### Perturbation and cNMF program interaction analysis

To evaluate associations between perturbations and cNMF programs, we fit a linear regression model using each cell’s log(program usage score+1) as the response and guide-positive status as the main predictor. Model covariates included inlet, log(nCount_RNA + 1), log(gRNA UMI count + 1), and percent mitochondrial reads. Standard errors were computed using the HC3 heteroskedasticity-consistent covariance estimator^110^. For emVar-program associations, NTC cells served as controls, and eligible cells carrying a guide targeting the corresponding variant element comprised the perturbation group, following the eligibility and variant-element mapping rules described above.

Association tests were performed for each perturbation state and each of the 30 cNMF programs. P-values were adjusted using the BH procedure separately within each program across all tested variant and positive-control perturbation states. Program-element associations with FDR < 0.10 were considered significant, and the direction of effect was determined from the sign of the regression coefficient. Fifteen programs showed at least one significant emVar-associated perturbation effect. P29 contained fewer than 30 genes meeting the |ρ| ≥ 0.1 threshold and was therefore excluded from figure-level characterization and subsequent analyses of emVar-associated programs, leaving 14 programs displayed in Figure 4B. Among the 14 retained programs, those significantly associated with perturbations of at least six distinct emVar loci at FDR < 0.10 were defined as convergent programs, and 85 significant emVar-program associations were identified.

#### Developmental trajectory analysis

To characterize the developmental expression of emVar-associated cNMF programs, we used two complementary human brain transcriptomic resources. First, program-gene expression was summarized across developmental age bins from the bulk cortical transcriptomic dataset^69^, and each program was standardized across developmental stages to define its temporal expression trajectory (Figure 4D). Second, for cellular-resolution analyses, we used single-cell transcriptomic data from the developing human prefrontal cortex^20^(Figure 4E, Figure S7C). For these analyses, we used the downsampled, normalized expression values provided by the single-cell transcriptomic data, which were designed to reduce differences attributable to sequencing depth across cell types and developmental stages.

For Figure 4E, we focused on five excitatory neuronal lineage clusters: L2-3_CUX2, L4_RORB, L5-6_THEMIS, L5-6_TLE4, and PN_dev. Program expression was calculated as the mean log-transformed CPM of program genes within each cell using the downsampled expression matrix and subsequently averaged across excitatory neuronal lineage cells within each of six developmental stages. Expression was then z-score-standardized within each program across the six stages. For Figure S7C, cell-type clusters were pooled into six broader lineage-level categories: astrocytes (Astro); excitatory neuronal lineage (L2-3_CUX2, L4_RORB, L5-6_THEMIS, L5-6_TLE4, and PN_dev); inhibitory neuronal lineage (VIP, ID2, LAMP5_NOS1, CGE_dev, SST, PV, PV_SCUBE3, and MGE_dev); oligodendrocyte lineage (OPC and Oligo); microglia (Micro); and vascular cells (Vas). Program expression was calculated from the same downsampled expression values, averaged within each lineage-developmental stage combination, and z-score-standardized within each program across the resulting 36 combinations. These analyses therefore reflect relative developmental and lineage-associated expression patterns within each program rather than absolute expression differences between programs.

#### Enrichment analysis of disease gene sets

External gene sets included ASD risk genes (Fu)^1^, DD risk genes (Kaplanis)^72^, NDDGeneHub Tier 1 and autosomal recessive genes and their ASD- and ADHD-associated subsets^70,73^, and gnomAD LoF-intolerant genes defined by pLI ≥ 0.9^75^ or LOEUF < 0.35^71^. ASD risk-gene sets (Fu) were evaluated at both FDR < 0.05 and FDR < 0.10. NDDGeneHub ASD and ADHD subsets were restricted to genes with non-empty annotations for the corresponding disorder. We also tested enrichment for ASD-versus-control DEGs at FDR < 0.05 across seven broad cell classes: astrocytes, blood–brain barrier cells, excitatory neurons, inhibitory neurons, microglia, oligodendrocytes, and oligodendrocyte precursor cells (Wamsley)^74^. Excitatory- and inhibitory-neuron results were displayed in Supplementary Figure S7D, and results for all seven classes are provided in Table S9.

To test whether each cNMF program gene set significantly overlapped these external gene sets, we performed one-sided Fisher’s exact tests. The background comprised 9,306 protein-coding genes eligible for mZINB association testing. For gnomAD, Fu, Kaplanis, and Wamsley gene sets, this background was restricted to genes evaluated in the corresponding resource. NDDGeneHub gene sets were tested against the full protein-coding background. For each gene, binary labels indicating membership in the program gene set and in the external gene set were used to construct a 2 × 2 contingency table. Fisher’s exact test P values were adjusted using the BH procedure separately across all 240 genetic-risk/constraint tests (30 programs × eight gene sets) and all 210 postmortem differential-expression tests (30 programs × seven cell classes). Enrichments with FDR < 0.10 were considered significant in the heatmaps and dot plots.

## KEY RESOURCES TABLE

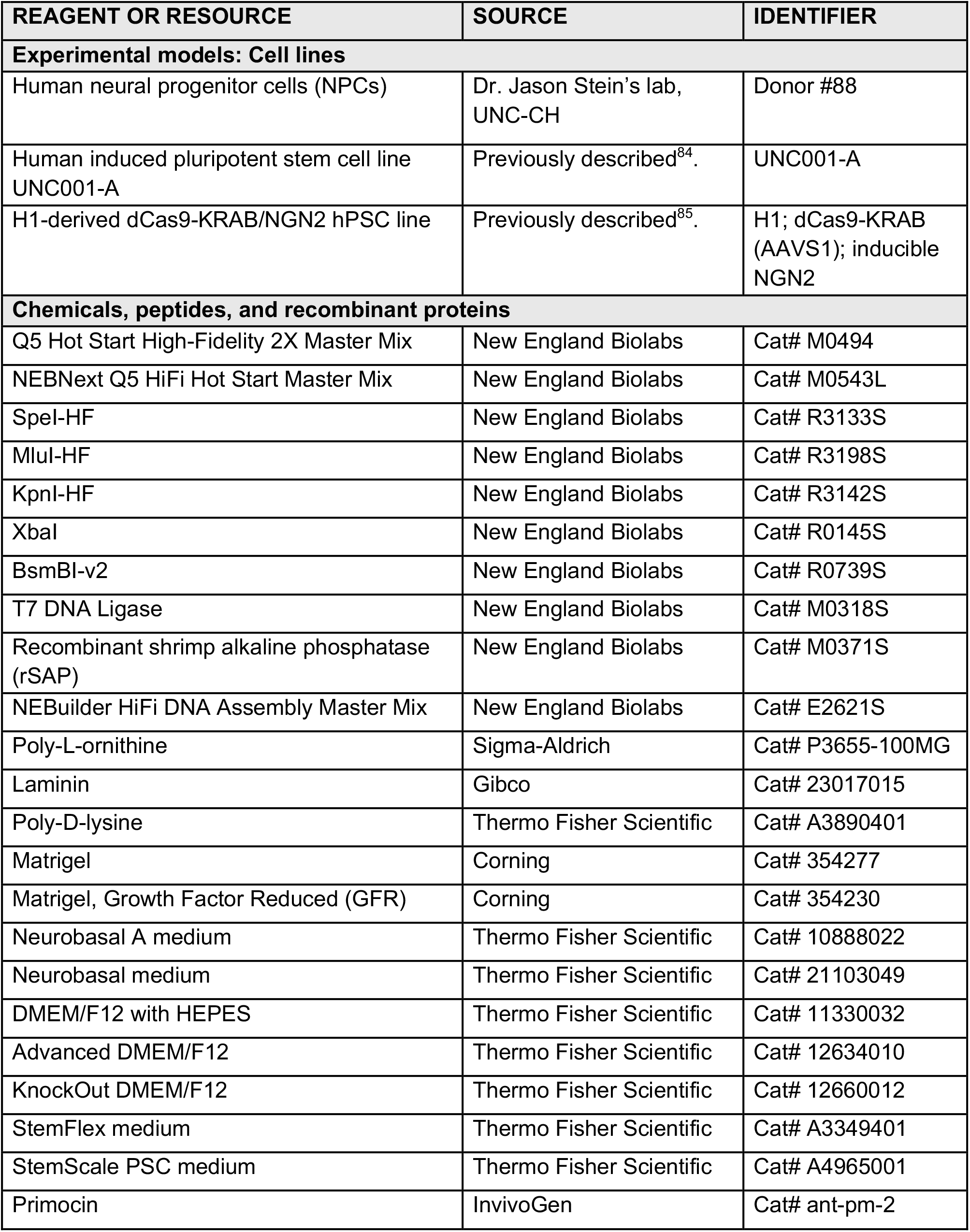
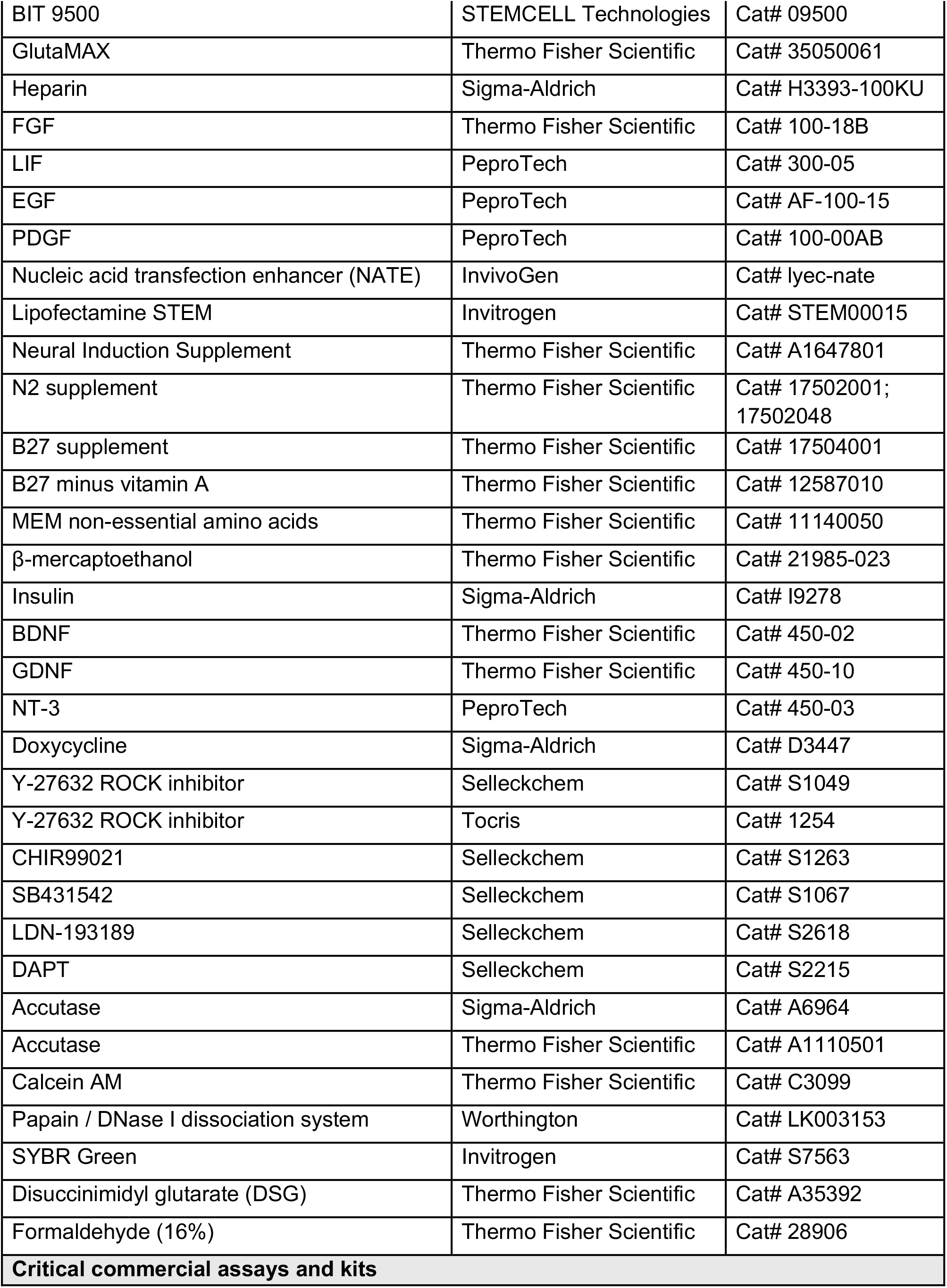
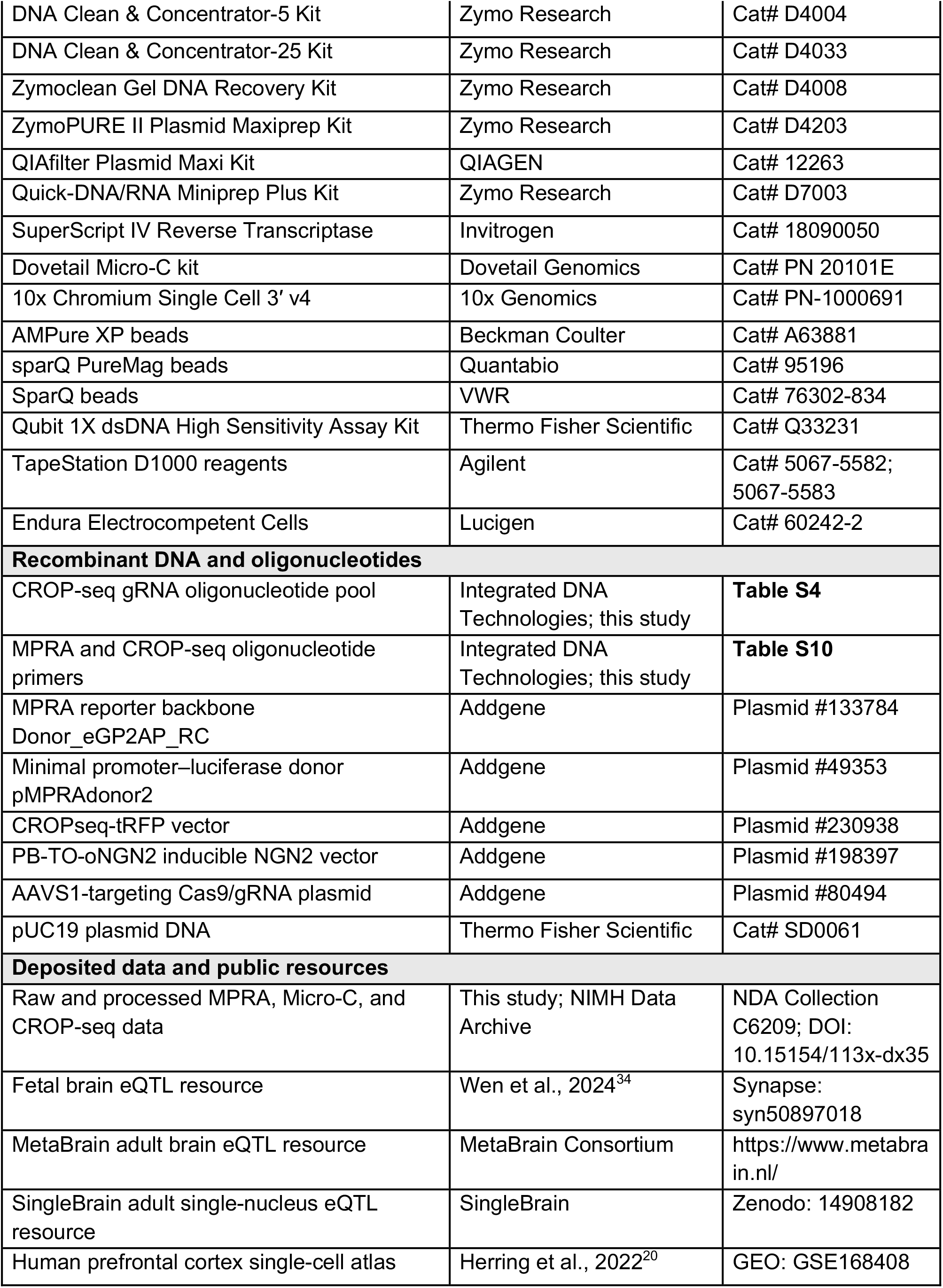
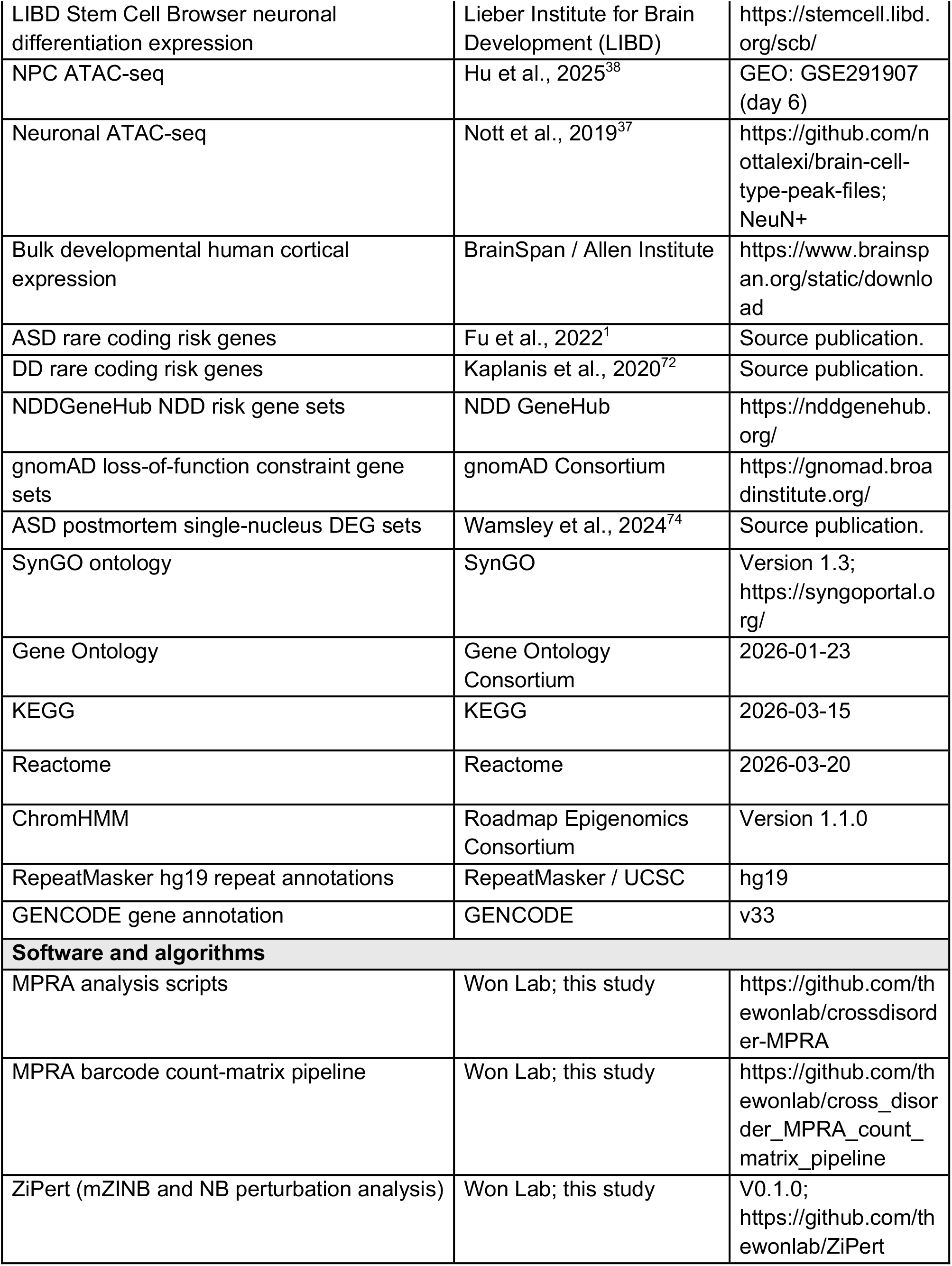
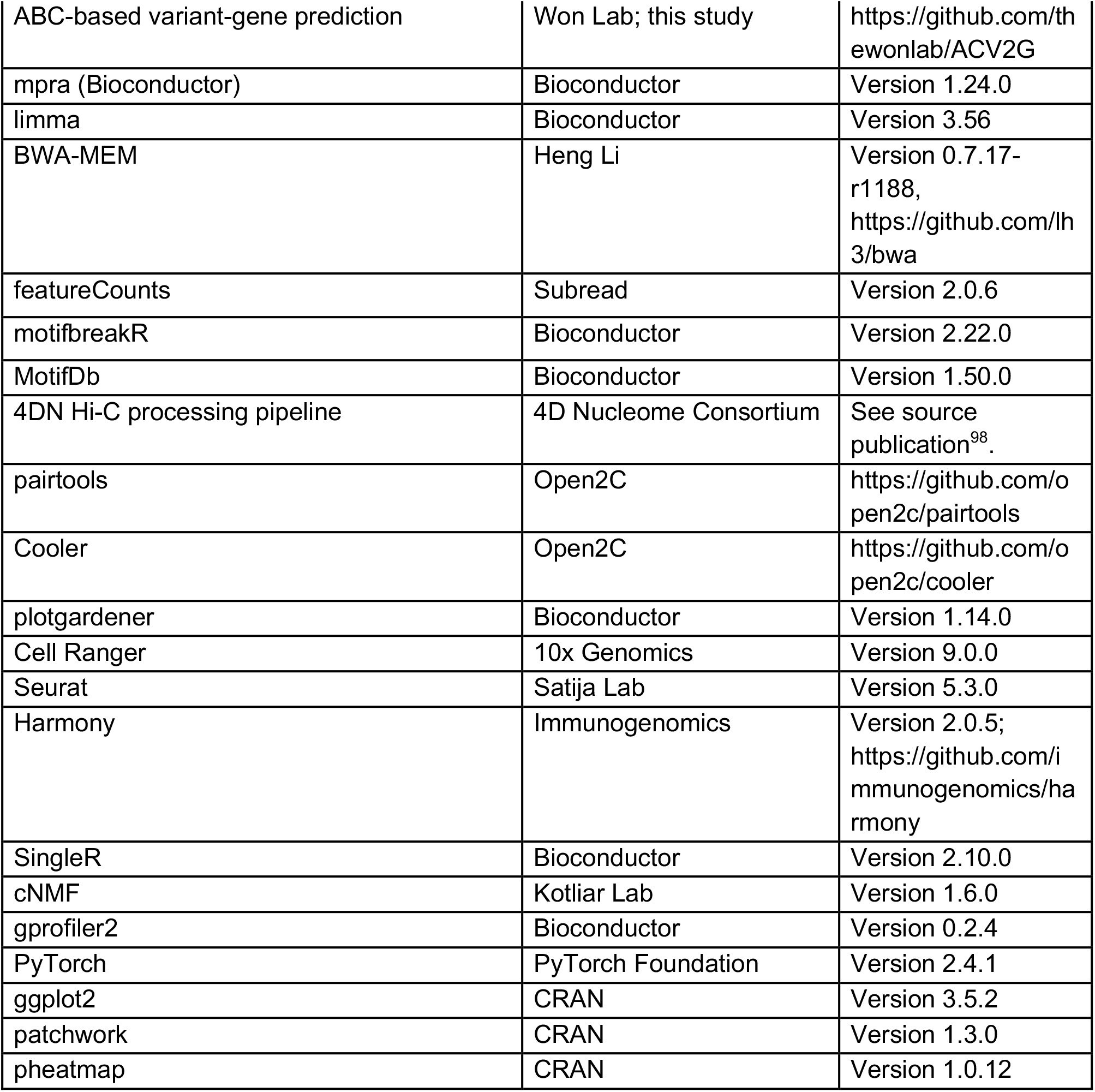

## SUPPLEMENTARY INFORMATION

### Supplementary Tables

**Table S1. MPRA summary statistics**

**Table S2. MotifbreakR transcription factor motif analysis**

**Table S3. Mapping emVars to genes via eQTLs and ABC model**

**Table S4. CROP-seq gRNA design and sequences**

**Table S5. CROP-seq analysis**

**Table S6. cNMF program gene membership and statistics**

**Table S7. cNMF program and emVar correlation analysis**

**Table S8. Functional enrichment analysis of emVar-dysregulated programs**

**Table S9. Neurodevelopmental disease risk gene set enrichment analysis**

**Table S10. MPRA and CROP-seq primer sequences**

### SUPPLEMENTARY FIGURES

**Supplementary Figure 1.**
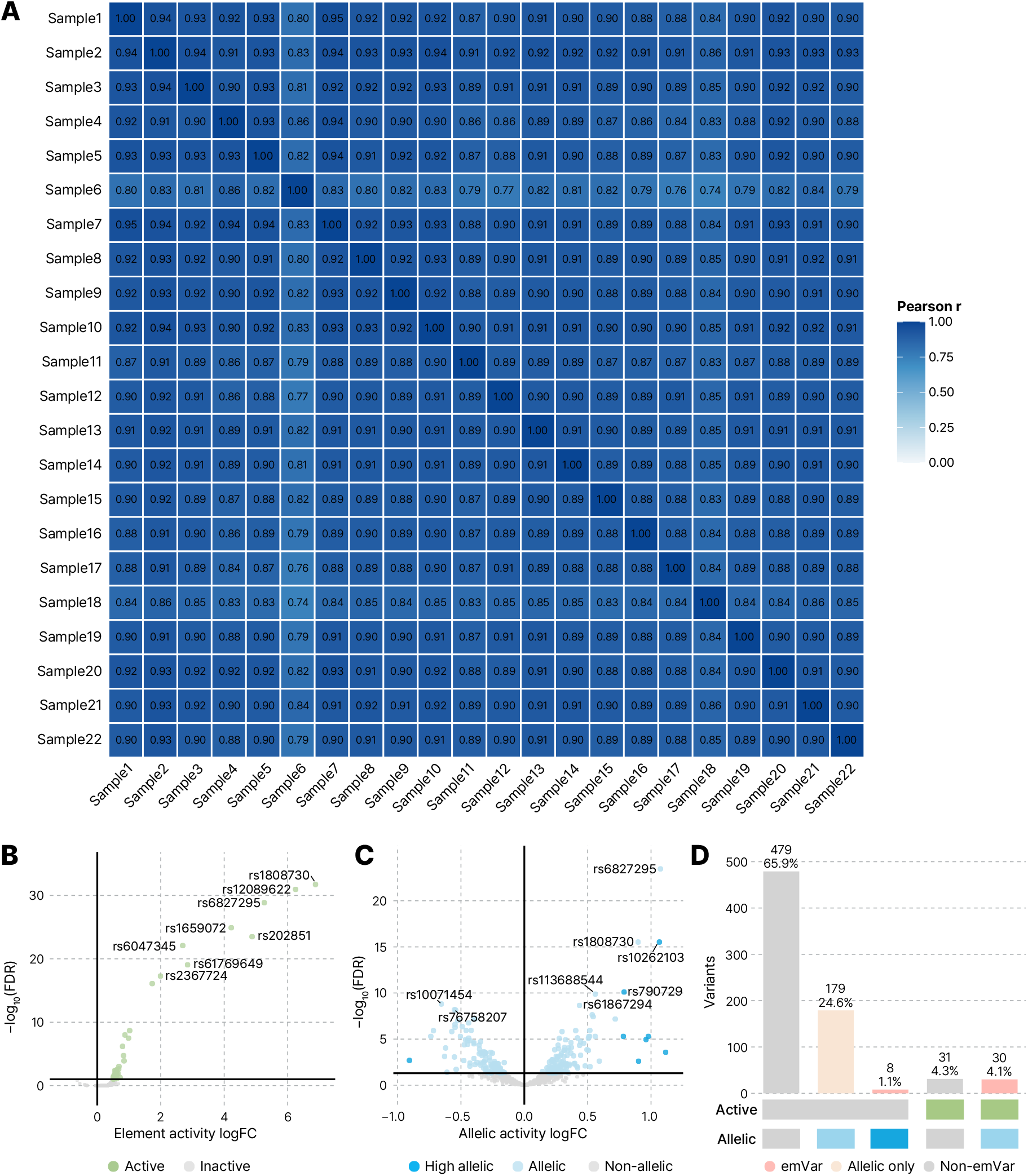
MPRA quality control and variant classification. **A.** Pairwise Pearson correlation heatmap of barcode-aggregated log(RNA/DNA) activity across 22 biological replicates (median Pearson r = 0.897 across 231 pairwise comparisons). **B.** Volcano plot of element-level MPRA activity. The x-axis shows element activity relative to randomly shuffled sequences (logFC), and the y-axis shows −log₁₀(FDR). Active elements, defined as FDR < 0.1 and logFC > 0, are highlighted in green. **C.** Volcano plot of allele-level MPRA activity. The x-axis shows allelic activity as logFC, and the y-axis shows −log₁₀(FDR). Variants with significant allelic activity (FDR < 0.05) are shown in light blue, and high-effect allelic emVars (|logFC| > 0.75) are shown in dark blue. **D.** Summary of MPRA regulatory categories across the 727 tested variants: active and allelic emVars (n = 30, 4.1%), high-effect allelic emVars (n = 8, 1.1%), active-only variants (n = 31, 4.3%), allelic-only variants (n = 179, 24.6%), and variants with neither active nor allelic activity (n = 479, 65.9%). The 38 variants classified as either active and allelic or high-effect allelic (pink) comprised the emVar set used for downstream analyses.

**Supplementary Figure 2.**
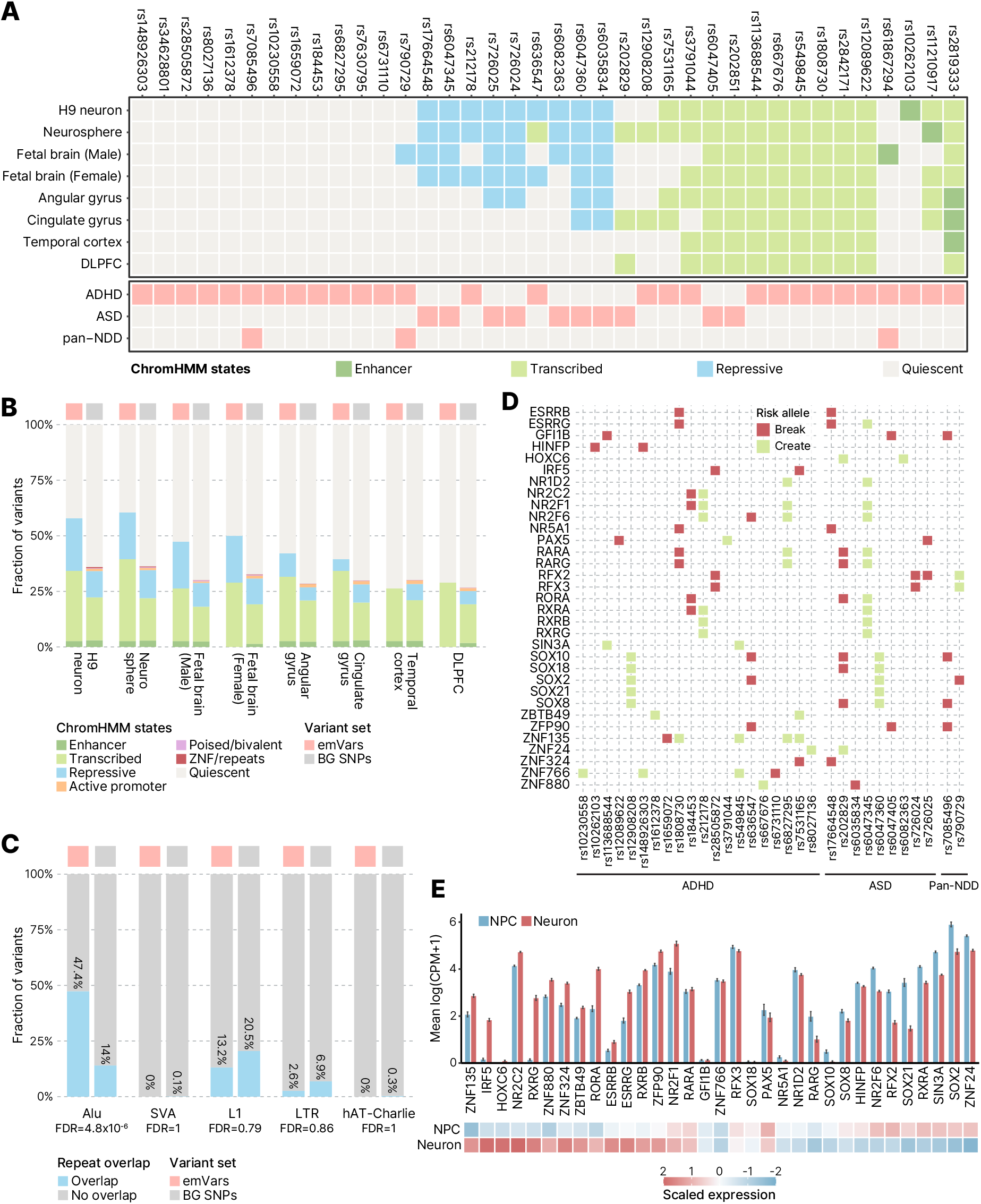
Genetic and epigenetic characterization of NDD emVars. **A.** ChromHMM state annotations for the 38 NDD emVars across hPSC-derived neurons (H9 neuron, neurosphere), fetal brains, and adult brains (angular gyrus, cingulate gyrus, temporal cortex, dorsolateral prefrontal cortex [DLPFC]). Columns represent individual emVars and rows represent cell or tissue contexts. Colors indicate broad ChromHMM state categories: enhancer (dark green), transcribed (light green), repressive (blue), and quiescent (gray). Bottom annotations indicate NDD association of each emVar (ADHD, ASD, or pan-NDD). **B.** Fraction of emVars and matched background SNPs assigned to each ChromHMM state across neural and brain contexts. Bars show the relative composition of enhancer, transcribed, repressive, active promoter, poised/bivalent, ZNF/repeats, and quiescent states. **C.** Overlap of emVars and matched background SNPs with selected repeat-element classes. Bars indicate the fraction of variants overlapping each repeat class. FDR values from Fisher’s exact tests with correction across the five repeat classes are shown for each comparison. **D.** Transcription factor binding motifs predicted to be disrupted (red) or created (green) by emVar risk alleles. Rows represent the 33 nominally enriched transcription factor motifs and columns represent individual emVars. Bottom annotations indicate the NDD GWAS association of each emVar. **E.** (Top) Mean expression of transcription factors represented in panel D in NPCs and neurons. Bars indicate mean log counts per million (CPM) + 1 expression. (Bottom) Gene-wise scaled expression of the same transcription factors across NPCs and neurons.

**Supplementary Figure 3.**
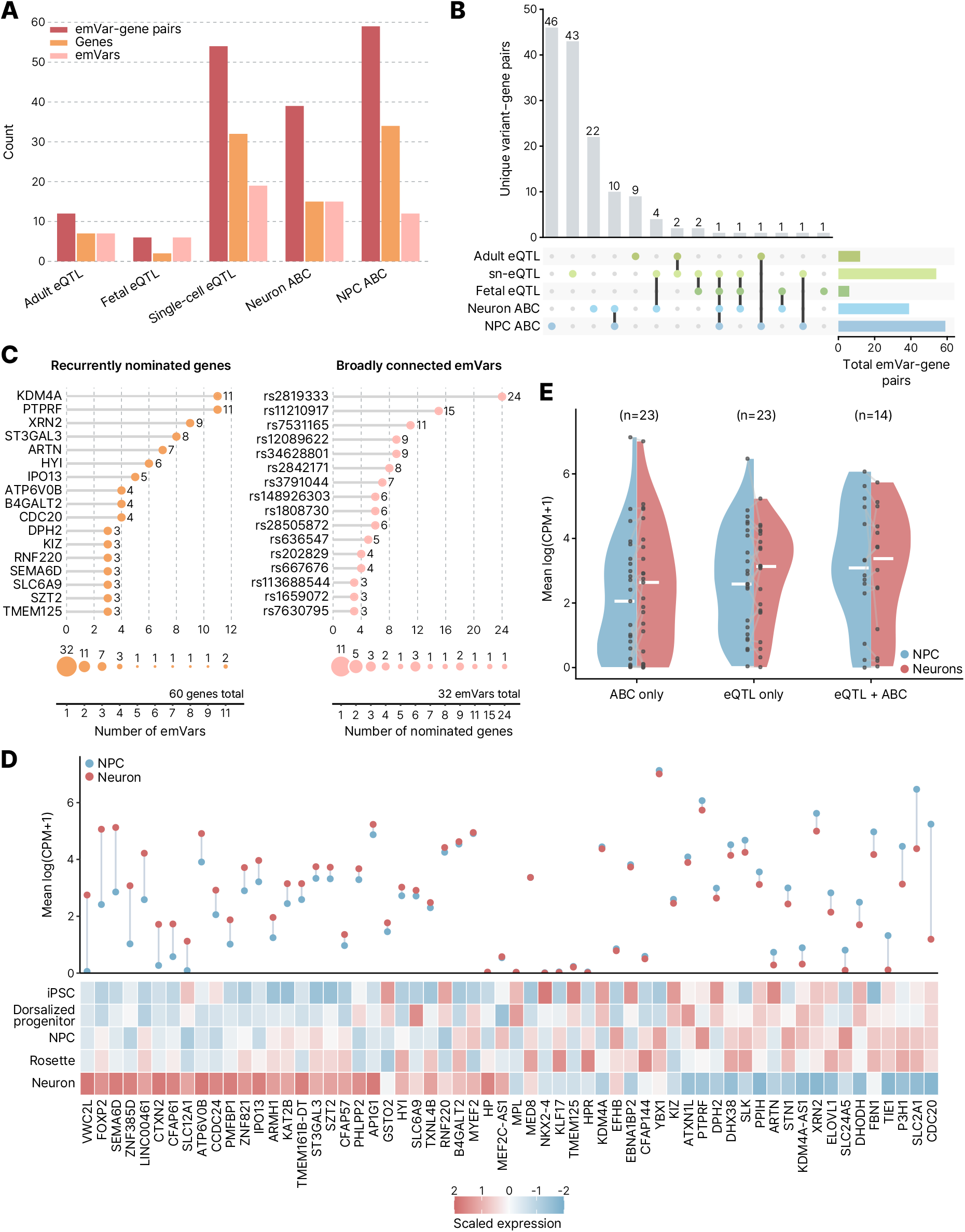
Comparison of emVar-gene mapping resources. **A.** Number of emVar-gene pairs, predicted target genes, and emVars nominated by each mapping resource. **B.** UpSet plot showing overlap of unique emVar-gene pairs across the five mapping resources. **C.** (Left) The upper plot shows genes nominated as targets of at least three emVars; points and adjacent numbers indicate the number of distinct emVars linked to each gene. The lower plot summarizes the complete distribution across all 60 nominated genes. Circle labels and sizes indicate the number of genes associated with the corresponding number of emVars. (Right) The upper plot shows emVars linked to at least three nominated target genes; points and adjacent numbers indicate the number of distinct genes linked to each emVar. The lower plot summarizes the complete distribution across all 32 emVars with at least one nominated target gene. Circle labels and sizes indicate the number of emVars associated with the corresponding number of nominated genes. **D.** Expression of emVar target genes across stem cell-derived neural differentiation states from the LIBD Stem Cell Browser. (Top) Mean log(CPM + 1) expression in NPCs and neurons, with lines connecting expression of the same gene across the two conditions. (Bottom) Heatmap showing gene-wise scaled expression across differentiation states. Columns represent nominated target genes, and rows represent differentiation states. **E.** Expression of nominated target genes in NPCs and neurons, stratified by gene-level mapping evidence. Each point represents a nominated gene, with lines connecting expression of the same gene between NPCs and neurons. Genes were classified as ABC only (n = 23), eQTL only (n = 23), or supported by both eQTL and ABC (n = 14). Violin plots show the distribution of mean log(CPM+1) expression within each group.

**Supplementary Figure 4.**
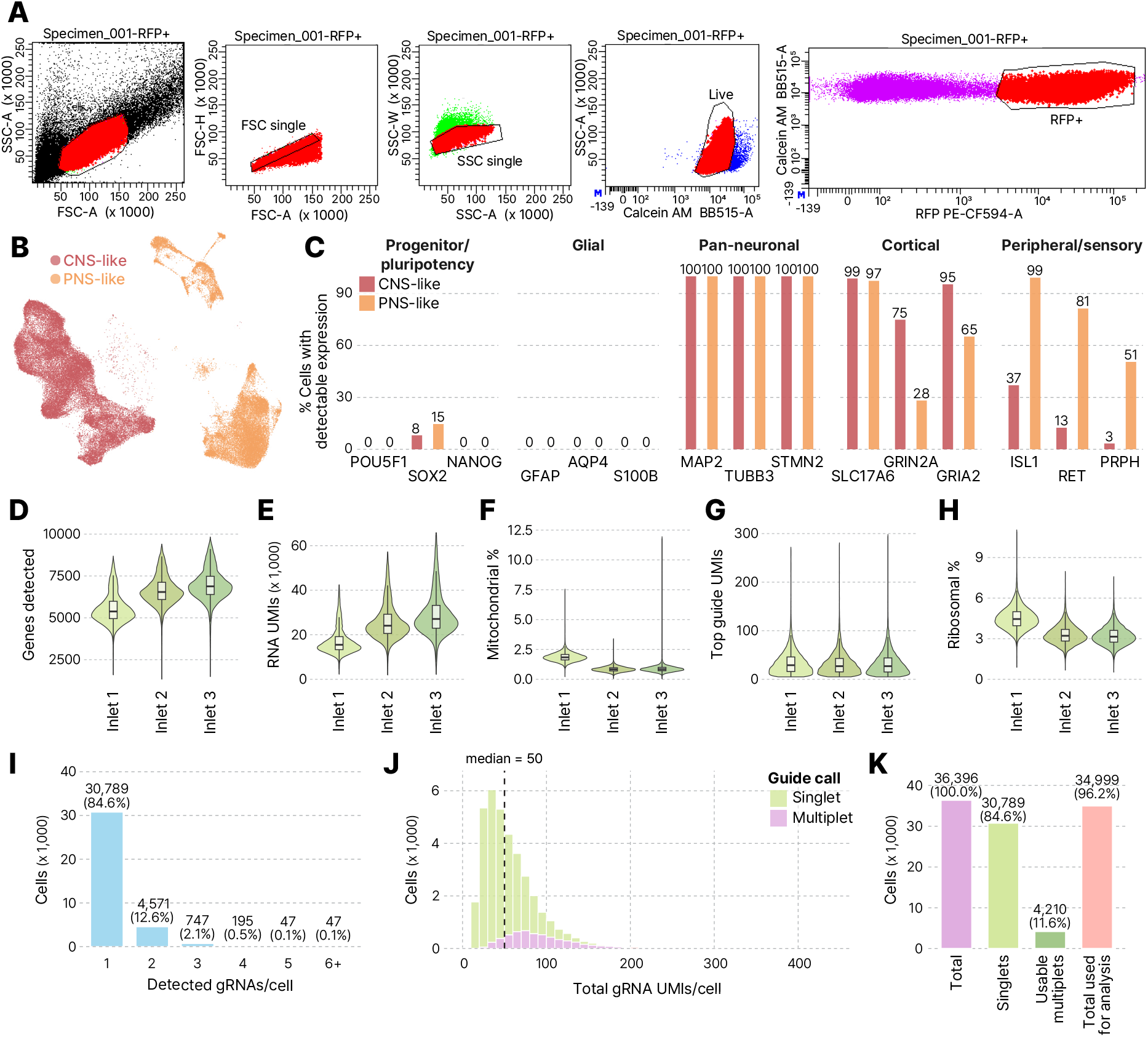
CROP-seq cell-level and gRNA assignment quality control for CNS-like neurons. **A.** Flow cytometry gating strategy used to isolate viable, gRNA-vector-positive cells for CROP-seq. Sequential gates excluded debris and doublets, selected live Calcein AM-positive cells, and enriched for RFP+ cells carrying the gRNA vector. **B.** Uniform Manifold Approximation and Projection (UMAP) visualization of QC-filtered neuronal cells annotated as CNS-like or PNS-like populations. PNS-like cells were excluded, and the CNS-like population was retained for downstream CROP-seq analyses. **C.** Percentage of CNS-like and PNS-like cells with detectable expression of selected progenitor/pluripotency (*POU5F1*, *SOX2*, *NANOG*), glial (*GFAP*, *AQP4*, *S100B*), pan-neuronal (*MAP2*, *TUBB3*, *STMN2*), cortical/excitatory-associated (*SLC17A6, GRIN2A*, *GRIA2*), and peripheral/sensory (*ISL1*, *RET*, *PRPH*) marker genes. **D-H.** Distribution of scRNA-seq and gRNA quality control metrics across the three sequencing inlets, including (D) number of genes detected per cell, (E) RNA UMI counts per cell, (F) percentage of mitochondrial reads, (G) top-guide UMI counts per cell, and (H) percentage of ribosomal reads. **I.** Number of detected gRNAs per cell. **J.** Distribution of total gRNA UMI counts per neuron, stratified as gRNA singlets (one detected gRNA) or multiplets (two or more detected gRNAs). The dashed line indicates the overall median of 50 gRNA UMIs per neuron. **K.** Summary of cells retained following cell-level and gRNA assignment quality control. Downstream CROP-seq analyses included gRNA singlets and multiplets meeting the inclusion criteria.

**Supplementary Figure 5.**
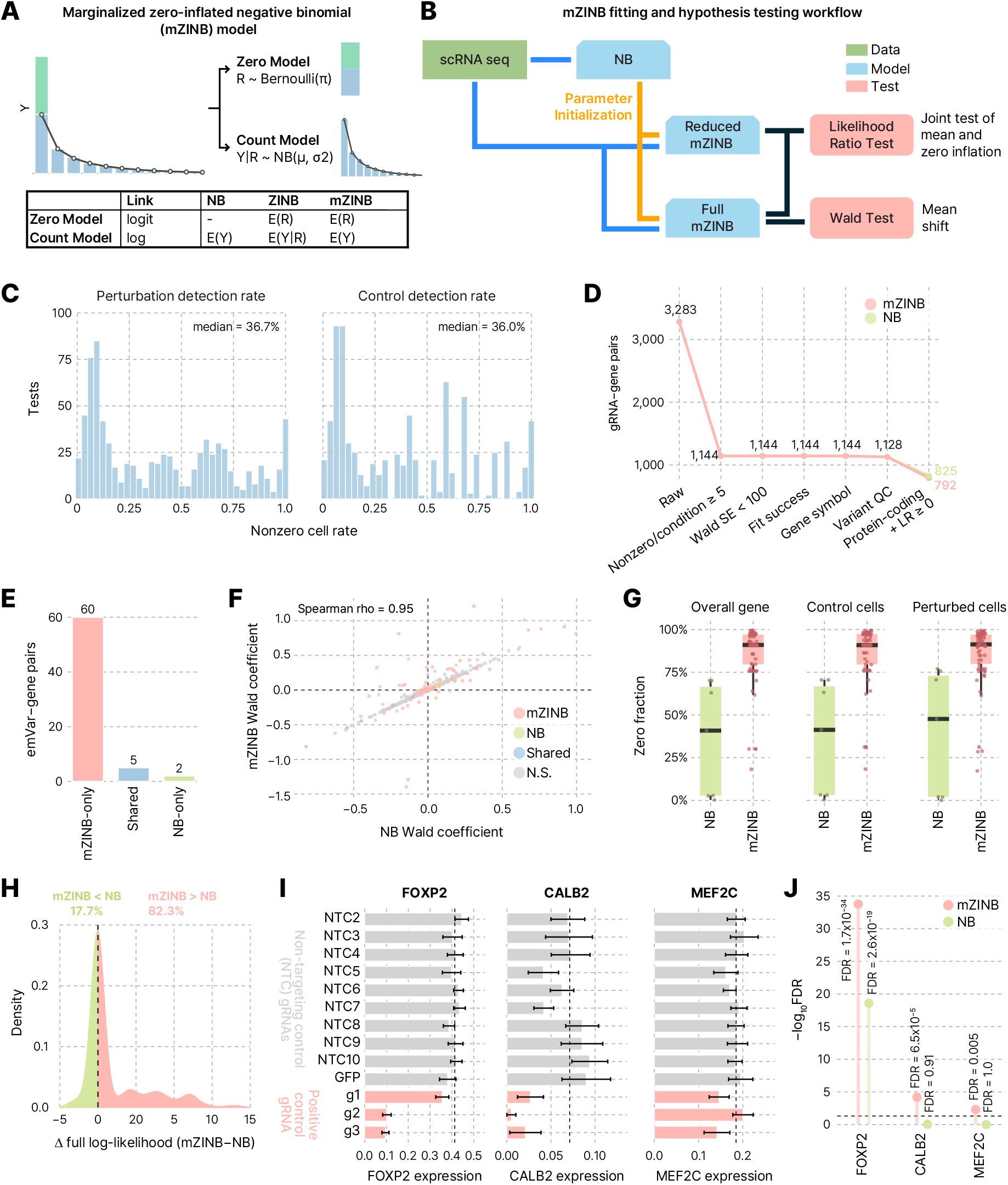
Sparsity-aware mZINB modeling, NB benchmarking, and positive control validation. **A.** Conceptual schematic of the mZINB model. The zero component models excess zero counts, while the negative binomial count component models gene-expression counts. The marginalized formulation estimates covariate effects on the overall mean expression scale while accounting for zero inflation. **B.** Schematic of the mZINB fitting and hypothesis-testing workflow. NB model estimates were used for parameter initialization of full and reduced mZINB models. The likelihood-ratio test (LRT) compares full and reduced mZINB models to jointly test perturbation-associated changes in mean expression and zero inflation, while the Wald test estimates the direction and magnitude of the mean-expression effect. **C.** Distribution of gene-detection rates among perturbation and control cells across candidate gRNA-gene tests prior to model fitting. **D.** Number of candidate gRNA-gene tests retained at each stage of model fitting and quality control for mZINB and NB. Both models were applied to the same 3,283 candidate tests and subjected to shared early filtering based on gene detection in at least 5 perturbed/control cells, Wald standard error, successful model fitting, gene annotation, and variant-level QC. These criteria retained 1,128 tests for both methods. Subsequent model-specific filtering for protein-coding genes and likelihood-ratio statistics (LR ≥ 0) retained 792 mZINB tests and 825 NB tests. **E.** Number of significant emVar-gene pairs identified uniquely by mZINB, by both models, or uniquely by NB. **F.** Concordance of Wald coefficients estimated by NB and mZINB across mZINB-only, NB-only, shared, and non-significant tests. Spearman correlation coefficient is shown. **G.** Zero-count fractions for genes identified as significant by NB or mZINB, calculated across all cells, control cells, and perturbed cells. **H.** Distribution of the difference in full-model log-likelihood between mZINB and NB. Positive values indicate higher log-likelihood under mZINB, whereas negative values indicate higher log-likelihood under NB. **I.** Mean log1p-transformed expression of *FOXP2*, *CALB2*, and *MEF2C* across cells assigned to individual NTC or positive-control gRNAs. Bars show mean expression, error bars indicate the standard error of the mean across cells, and dashed lines indicate the corresponding mean expression across NTC gRNAs. **J.** Significance of *FOXP2*, *CALB2*, and *MEF2C* positive-control perturbations under mZINB and NB. Dashed horizontal line indicates FDR = 0.05.

**Supplementary Figure 6.**
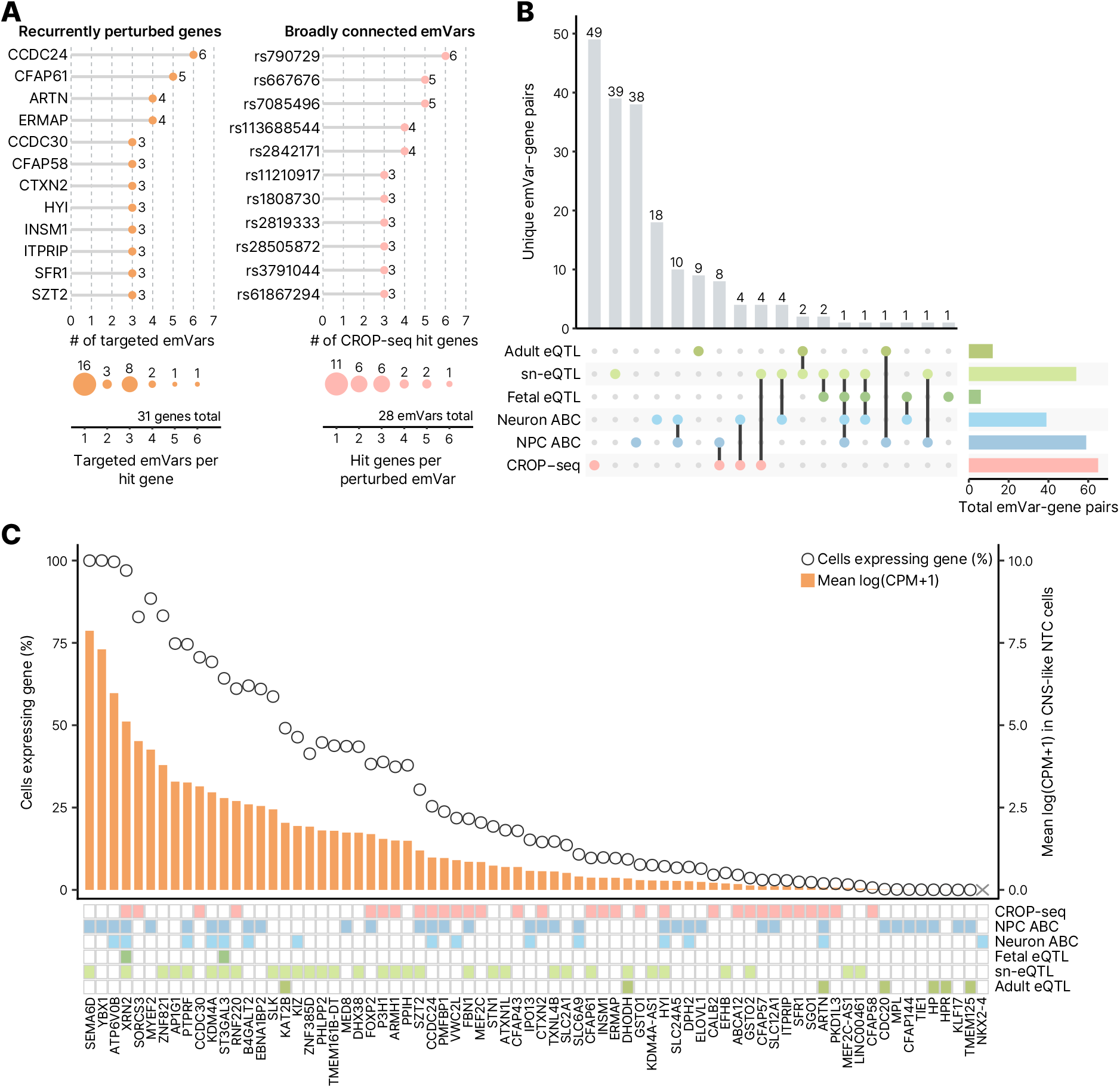
Comparison of CROP-seq-supported and predicted emVar-gene relationships across mapping sources. **A.** (Left) The upper lollipop plot shows recurrently perturbed genes, defined as genes significantly associated with at least three targeted emVars. Points and adjacent numbers indicate the number of distinct emVars associated with each gene. The lower dot plot summarizes the distribution across all 31 CROP-seq hit genes, with dot size and labels indicating the number of genes associated with each number of targeted emVars. (Right) The upper lollipop plot shows broadly connected emVars, defined as emVars significantly associated with at least three CROP-seq hit genes. Points and adjacent numbers indicate the number of distinct hit genes associated with each emVar. The lower dot plot summarizes the distribution across all 28 emVars with at least one significant CROP-seq association, with dot size and labels indicating the number of emVars associated with each number of hit genes. **B**. UpSet plot showing the overlap of unique emVar-gene pairs identified by adult brain eQTL, fetal brain eQTL, sn-eQTL, neuron-ABC, NPC-ABC, and CROP-seq. Vertical bars show the number of pairs unique to each indicated resource combination, connected dots identify the resources contributing to each intersection, and horizontal bars show the total number of pairs contributed by each resource. **C.** Baseline expression and resource membership of the 74 genes represented in the union of the six mapping resources. Orange bars show mean log(CPM+1), and open circles show the percentage of cells with detectable expression, calculated across 5,920 cortical/CNS-like NTC cells from the filtered CROP-seq dataset. Genes are ordered by decreasing mean expression from left to right. The annotation matrix below indicates whether each gene participated in at least one emVar-gene pair identified by the corresponding resource. The gray cross at *NKX2-4* denotes that no gene expression counts were detected in the filtered CROP-seq gene expression matrix.

**Supplementary Figure 7.**
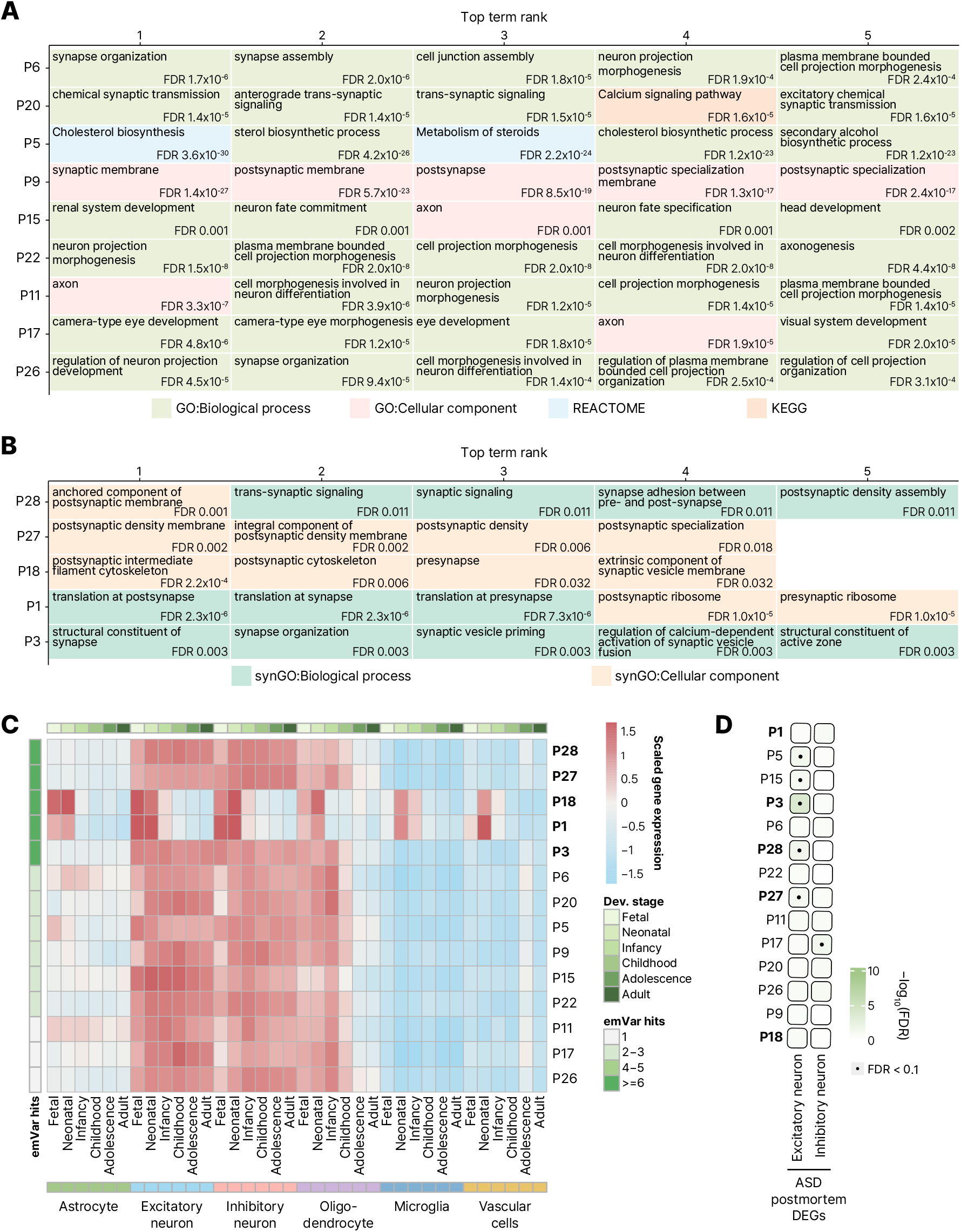
Functional and developmental characterization of cNMF programs. **A.** Top five FDR-significant GO, KEGG, or Reactome terms for each of the nine retained non-convergent programs. Terms are ranked by FDR, with rank 1 indicating the most significant term. **B.** Top SynGO terms for the five convergent programs. Terms are ranked by FDR, with rank 1 indicating the most significant term. Colors in A and B indicate the corresponding annotation database or ontology category. **C.** Developmental and cell type lineage-level expression profiles of the 14 retained cNMF programs across a human brain single-cell atlas. Rows represent programs, and columns represent broad neural or glial lineage-developmental stage combinations. Colors indicate within-program z-scored expression across the 36 cell type-developmental stage combinations. The five convergent programs are shown in bold at the top. Dev. stage, developmental stage. **D.** Enrichment of the 14 retained cNMF programs for DEGs identified in excitatory and inhibitory neurons from ASD postmortem brains. Color intensity indicates –log10FDR, and dots indicate enrichments with FDR < 0.1 based on Fisher’s exact test. The five convergent programs are shown in bold.

## Notes

### Competing Interest Statement

The authors have declared no competing interest.

